# A Graph-based QSAR Modeling Pipeline for Predicting In vitro PubChem Assays and In vivo Human hepatotoxicity: Mechanistic Analysis of Caspase-3/7 Activation

**DOI:** 10.64898/2026.06.10.731399

**Authors:** Yamini Chitikela, Zhenquan Jia, Chunjiang Zhu

## Abstract

Caspase-3 and -7 are key effector caspases in the apoptotic pathway, a form of programmed cell death, and their activities serve as a well-established biomarker for evaluating environmental chemical toxicity and informing chemical risk assessment. Loss of mitochondrial membrane potential is a key event in the activation of Caspase-3/7 signaling and the subsequent induction of apoptosis. Therefore, simultaneous assessment of mitochondrial membrane potential and Caspase-3/7 activity enables elucidation of the mechanisms and pathways through which apoptosis is initiated. Quantitative Structure–Activity Relationship (QSAR) modeling have been widely used for toxicity prediction. While advanced graph deep learning approaches, such as graph transformers (GTs), have shown outstanding performance in many domains, they have not been fully leveraged in QSAR modeling. We developed a novel graph-based Quantitative Structure–Activity Relationship (QSAR) modeling pipeline that has not previously reported, encompassing assay data preprocessing, feature representations (fingerprints and molecular graphs), and benchmarking machine learning (ML) models, including classic ML models, graph neural networks, graph transformers, and their consensus ensembles. The pipeline has been applied to predict environmental chemical-induced Caspase-3/7 activation, mitochondrial membrane potential disruption, and FDA-approved Drug-Induced Liver Injury (DILI), enabling a sensitive and mechanism-informed framework for assessing human hepatotoxicity. For example in the prediction of FDA-approved DILI, the graph transformer method, Graphormer, had the highest F1 score 0.79 and the full consensus model achieved the highest AUC 0.69, significantly improving the previous best model with AUC 0.63 and F1 0.65. We also investigated cell-line-specific responses to environmental chemical exposure by identifying structural motifs that selectively induce Caspase-3/7 activation in individual cell lines. Human embryonic kidney (HEK293) was the only cell line with a distinct structural motif: 1,1-dichloroethane and chlorobenzene. Human neuroblastoma (SK-N-SH) is uniquely impacted by an epoxide fragment.

**For Table of Contents Only:** 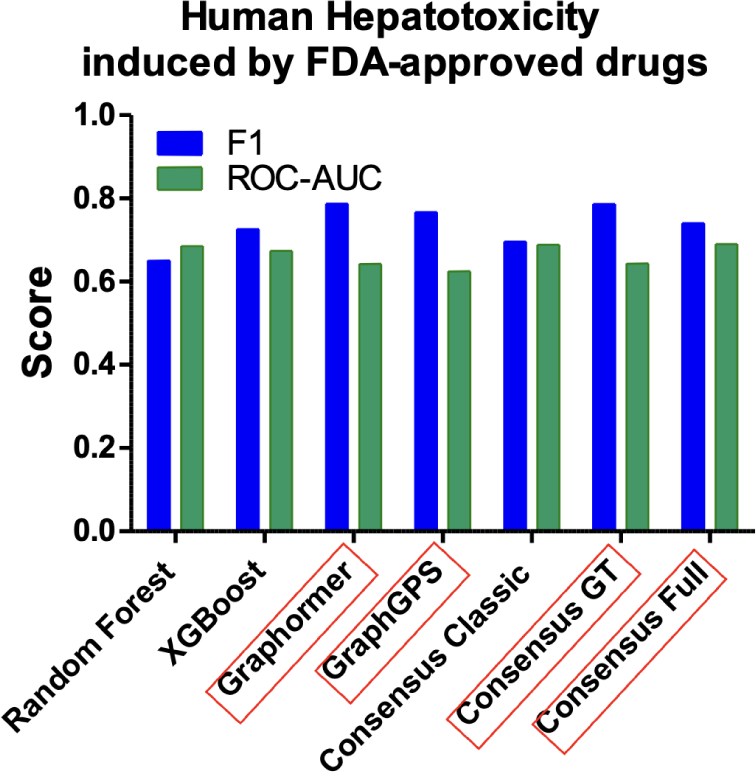

## Introduction

Apoptosis is a form of programmed cell death mediated by Caspase-3/7 that is crucial for maintaining internal homeostasis [1]. As the central enzymes of the apoptotic execution phase, Caspase-3/7 constitute the common pathway through which cells irreversibly commit to apoptosis. Environmental compounds can induce excessive apoptosis by activating Caspase-3/7; therefore, monitoring their activity serves as a key indicator for assessing the apoptotic toxicity of such compounds. The collapse of mitochondrial membrane potential is an early event in intrinsic apoptosis that triggers the Caspase cascade [2]. The *simultaneous* monitoring of membrane potential and Caspase-3/7 activity enables the systematic capture of the entire process from mitochondrial damage to the execution of apoptosis, thereby providing mechanistically defined biomarkers for evaluating the apoptotic toxicity of compounds.

Rapid and accurate assessment of the potential toxicity of environmental chemicals and drugs remains a major challenge facing the field. Computational predictive models can serve as a powerful complement to traditional experimental methods, aiding in the prioritization and strategic direction of toxicity testing. Consequently, such models hold the promise of not only shortening testing cycles and reducing costs but also minimizing or even eliminating reliance on animal experimentation. Quantitative high-throughput screening (qHTS) generates full concentration–response curves for each compound, reducing false positives and false negatives relative to single-dose assays [3]. Xia *et al.* [4] profiled 1,408 compounds for cytotoxicity across 13 human and rodent cell lines, showing that no single cell line can serve as a universal surrogate for organ toxicity. The Toxicology in the 21st Century (Tox21) programme subsequently scaled these assays to 10K compounds using an automated robotic platform [5]. The Tox21 data, publicly available in PubChem BioAssay [6], has greatly enabled the development of computational modeling for predicting *in vivo* toxicity to replace costly and time-consuming animal testing.

Quantitative Structure–Activity Relationship (QSAR) modeling has been a cornerstone of computational toxicology since Hansch and Fujita [7]. QSAR models have been widely used for toxicity prediction [8]. Models trained on the full Tox21 10K profiles demonstrated that a combination of structural and activity data results in better models than using structure or activity data alone [9]. Deep learning algorithms have been shown to outperform classic random forest and support vector machine in Tox21 assays [10]. For example, Kim *et al.* [11] studied hepatotoxicity and developed a QSAR model based on the antioxidant response element beta lactamase reporter gene assay from the Tox21. However, studies specifically targeting *Caspase-3/7 activation*, a key indicator of apoptosis, remain relatively scarce.

To date, only one such report has been published, conducted by Huang *et al.* [12], who analyzed Caspase-3/7 activation and cell viability data in Tox21 and proposed a fragment-based weighted feature significance (WFS) QSAR model. This model achieved performance comparable to Naive Bayesian models and superior to SVM, while offering improved interpretability through feature-level analysis [12]. The WFS model proposed in this study constructs a simple yet highly interpretable framework for toxicity prediction by statistically quantifying the enrichment of “toxic structural features” within active compounds. Unlike methods relying on overall molecular similarity, this approach is grounded in fragment-level structural information, thereby enabling the effective prediction of toxicity for structurally diverse compounds. However, the model still presents key limitations. Its training dataset integrates high-throughput screening results derived from 13 distinct cell lines, defining “pan-cytotoxicity” through averaging or aggregation without explicitly distinguishing the biological differences among these various cell types during the modeling process. This approach implicitly assumes that the toxic effects of a compound exhibit uniformity across different cell types. However, different cells demonstrate significant variations in metabolic capacity, receptor expression (particularly upstream signaling receptors associated with Caspase-3/7 activation), and signaling pathways. Consequently, the same compound may induce distinct or even opposing toxic responses in different cellular contexts, including differential effects on cell viability and Caspase-3/7 activation.

Furthermore, the quality of QSAR models depends on the machine learning approach, the molecular descriptors used for the toxicity endpoint, and the quality of the training data [12]. Prior work using Tox21 qHTS data has demonstrated that models combining chemical fingerprints with bioactivity data achieve higher predictive accuracy than structure-only models, particularly for endpoints with clear mechanistic interpretations [9, 10]. However, classical fingerprint-based approaches encode molecular topology as fixed-length binary vectors and may lose structural information. **Graph-based approaches** encode compounds directly as molecular graphs with atoms as nodes and bonds as edges, allowing structure-activity relationships to be learned from molecular topology without the information loss inherent in binary fingerprints [13]. Classical QSAR fingerprints encode molecules as fixed-length binary vectors, discarding bond connectivity [14]. *Graph neural networks (GNNs)* encode each compound as a molecular graph, learning representations through iterative message passing [13]. Multiple GNN architectures—GCN, GAT, GIN, and GraphSAGE—have been proposed with different aggregation strategies.

The attention-based transformer architecture has demonstrated outstanding predictive performance across multiple domains [15], such as topping the leaderboard in the molecular property prediction [16]. As a key technique for the emerging large language models (LLMs), it has been fundamental for the powerful prediction capability of LLMs. Inspired by the success of transformers in natural language processing, graph transformer research applies transformers in graph data through tokenization and structural and positional encoding techniques. *Graph transformers (GTs)* aggregate information from all nodes in a given graph, thus not suffering local structural bias in GNNs [17]. However, **graph transformers (GTs) have not been fully leveraged in QSAR modeling on in vitro assay modeling and in vivo human hepatoxocity. To the best of our knowledge, there is no prior QSAR study on a comprehensive evaluation of fingerprint-based and graph-based approaches (GNNs and GTs) for Caspase and mitochondrial toxicity prediction**.

Moreover, applicability domain analysis has been established best practices for QSAR model development [8]. The applicability domain defines the chemical space in which model predictions are considered reliable; compounds outside this region should be treated with caution [18].

In this study, we propose a graph-based QSAR modeling pipeline that encompasses assay data preprocessing, feature representations (fingerprints and molecular graphs), and benchmarking machine learning (ML) models, including classic ML models, GNNs, advanced graph transformers, and their consensus ensembles. The developed pipeline enabled a comprehensive study on the performance of various machine learning models as QSAR models, including three classical ML classifiers, four GNNs, two GTs, and four consescus models. Based on *in vitro* Caspase and mitochondrial assays in PubChem, we applied the pipeline to predict Caspase-3/7 activation and mitochondrial membrane potential disruption. Experimental evaluations show that GTs and GNNs outperformed classic ML models when the number of active compounds is large, such as MMP disruption, while GTs and classic ML models performed good for highly imbalance data with limited active compounds, such as Caspase-3/7 activation. Since these QSAR models for predicting toxicity on *in vitro* assays usually do not generalize well to *in vivo* human hepatotoxicity [18], we also conducted graph-based QSAR modeling for FDA-approved Drug-Induced Liver Injury (DILI) dataset [19]. Here, the full consensus model achieved the highest AUC 0.69 and Graphormer had the highest F1 score 0.79, both surpassing the previous best model with AUC 0.63 and F1 0.65 with a large margin.

Moreover, the interplay between mitochondrial dysfunction and Caspase-3/7 activation remains unexplored, representing a critical gap in linking early mechanistic events to downstream toxicological outcomes. Next, we performed a cross-assay mechanistic analysis to characterize the relationship between mitochondrial perturbation, Caspase-3/7 activation, and cell viability, providing insight into the apoptotic mechanisms induced by tested compounds. We conducted a comprehensive comparison between MMP disruption pathway and Caspase-3/7 activation to identify chemical substructures that are presented in compounds with dual activations. For example, phenolic compounds bearing a para-hydroxyphenyl motif, as well as members of the lipophilic chain family with long alkyl chains can trigger the collapse of MMP, leading to the activation of Caspases-3 and -7. To understand the impacts of different cellular contexts on toxic responses for the same compound, we identified *cell-line-specific* chemical substructures that exclusively activate Caspase-3/7 in a specific cell line but not in all other cell lines. Human embryonic kidney (HEK293) was the only cell line with a distinct structural motif: 1,1-dichloroethane and chlorobenzene. Human neuroblastoma (SK-N-SH) is uniquely impacted by an epoxide fragment and rat hepatoma (H-4-II-E) is uniquely impacted by a tetramethylcyclohexene motif and an acetaldehyde fragment. We also offered fine-grained insights into the interplay with MMP disruption and cell viability, particularly compounds with different activation outcomes in MMP and cell viability assays.

## Methodology

**Figure 1** shows an overview of our proposed QSAR modeling pipeline which we use for predicting Caspase-3/7 activation, mitochondrial membrane potential, and FDA DILI gold standard, and we explain the details in this section.

**Figure 1:**
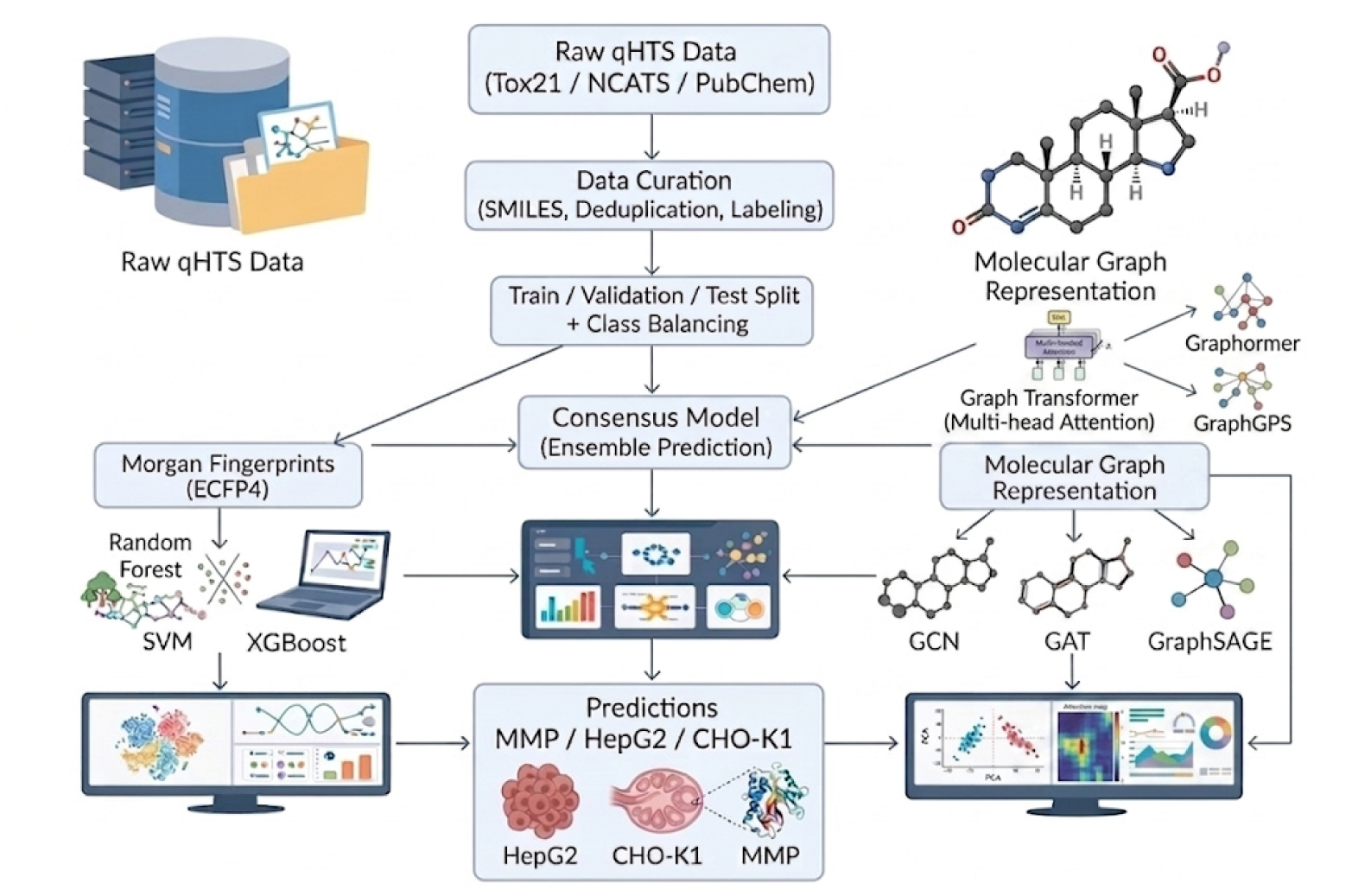
Overview of the QSAR modeling pipeline integrating data preprocessing, feature representations, and machine learning models from classic machine learning and graph-based approaches.

### Databases: Sources and Preprocessing

PubChem is a database of chemical molecules and their activities against biological assays, managed by the US National Institutes of Health. We downloaded from PubChem six assays from the Tox21 database and thirteen assays from the National Centre for Advancing Translational Sciences (NCATS), all in the CSV format. **Table 1** summarizes brief introductions on the different assays, their PubChem AID, and total number of compounds.

**Table 1:** Summary of all nineteen qHTS assays used in this study. MMP = mitochondrial membrane potential; Viab. = cell-viability counter screen.

| Cell Line | Description | Total | AID |
| --- | --- | --- | --- |
| *NCATS Caspase-3/7 Assays* |  |  |  |
| HepG2 | Human liver carcinoma | 1,408 | 654 |
| Jurkat | Human T-cell leukaemia | 1,408 | 655 |
| HUV-EC-C | Human vascular endothelial | 1,408 | 656 |
| SH-SY5Y | Human neuroblastoma | 1,408 | 657 |
| BJ | Normal human fibroblast | 1,408 | 658 |
| MRC-5 | Normal human lung fibroblast | 1,408 | 659 |
| Mesangial | Mouse glomerular mesangial | 1,408 | 660 |
| SK-N-SH | Human neuroblastoma | 1,408 | 661 |
| H-4-II-E | Rat hepatoma | 1,408 | 663 |
| HEK293 | Human embryonic kidney | 1,407 | 664 |
| N2a | Mouse neuroblastoma | 1,408 | 665 |
| NIH 3T3 | Mouse embryonic fibroblast | 1,408 | 666 |
| Renal Proximal Tubule | Human renal epithelial | 1,408 | 667 |
| *Tox21 Caspase-3/7 Assays* |  |  |  |
| HepG2 | Human liver carcinoma | 9,667 | 1346978 |
| HepG2 | Human liver carcinoma (Viab.) | 9,667 | 1346981 |
| CHO-K1 | Chinese hamster ovary | 9,667 | 1346980 |
| CHO-K1 | Chinese hamster ovary (Viab.) | 9,667 | 1346979 |
| *Tox21 MMP Assays* |  |  |  |
| HepG2 | Human liver carcinoma (MMP) | 10,486 | 720635 |
| HepG2 | Human liver carcinoma (MMP Viab.) | 10,486 | 720634 |

From Tox21, we use the qHTS assay for Caspase-3/7 induction by small molecules in HepG2 cells (AID_1346978) and the cell-viability counter screen (AID_1346981). HepG2 is a human hepatocellular carcinoma cell line widely used to model hepatotoxicity.

We also use a qHTS assay for Caspase-3/7 induction in CHO-K1 cells (AID_1346980) and its cell-viability counter screen (AID_1346979). CHO-K1 is a subclone of the parental Chinese hamster ovary (CHO) cell line derived from the ovary of an adult female Chinese hamster; it is widely employed in toxicology research as a *non-human* mammalian reference cell line, providing a complementary species perspective to the human HepG2 data [20].

In addition, we adopt thirteen NCATS assays measure Caspase-3/7 cellular toxicity across thirteen distinct cell lines, spanning a broad range of tissue types and organisms. Combining with Tox21 assays, fourtheen cell types are considered in this study. Table 1 provides a short summary and detailed descriptions on diverse cell lines are referred to Supplementary S1.

We use a qHTS assay for small molecule disruptors of the MMP in HepG2 cells (AID_720635). In addition to the mitochondrial potential assay, these cells were also used to test the cytotoxicity by determining the cell viability in the culture based on the quantification of ATP present (a crucial molecule for various cellular processes). The cytotoxicity (AID_720634) was measured in the same assay well of the MMP assay.

#### Activity Score and Outcome

Each assay file includes a continuous activity score derived from the concentration-response curve fitting. When dose-response data are available, the Fit_LogAC50 value is used to quantify potency: it represents the logarithm (base 10) of the half-maximal activity concentration (AC50) in molar units, estimated by fitting a four-parameter Hill equation to the observed response curve. This value is then used to calculate a relative activity score by mapping the Fit_LogAC50 onto a scaled score range defined for each curve class, such that more potent compounds (lower AC50 values) receive higher scores. The resulting scores are normalised to the range of 0 to 100.

Activity classification follows PubChem conventions based on the computed activity score. Compounds with a score of 0 are classified as *inactive*; those with scores between 40 and 100 are classified as *active*; and those with scores between 1 and 39 are classified as *inconclusive*. The inconclusive category captures compounds with intermediate concentration-response behaviour that cannot be reliably assigned to either outcome.

#### Data Cleaning

Before analysis, metadata rows were removed from each downloaded assay file. Duplicate records were then resolved by sorting all records for each assay by compound ID (PubChem_CID) and activity score (PUBCHEM_ACTIVITY_SCORE) in descending order of score, then retaining only the first record per compound. This ensures that when a compound appears multiple times, whether with the same or different activity outcomes, only the single record with the *highest* activity score is kept, preventing underestimation of compound activity. Keeping only one record for each compound also facilitates AI model training with a unique outcome.

**Table 2** illustrates the procedure with two representative cases. In Case 1, the compound with CID 1215403 appears twice with the same Inconclusive outcome but different scores (10 and 3); the higher-scoring record is retained. In Case 2, the compound with CID 9290403 appears three times in two different outcomes: Active (scores 45 and 22) and Inactive (score 0). Only the record with the highest overall score—Active, score 45—is retained; all others are discarded. Note that the same cleaning procedure was applied to all Tox21 and NCATS assay files.

**Table 2:**
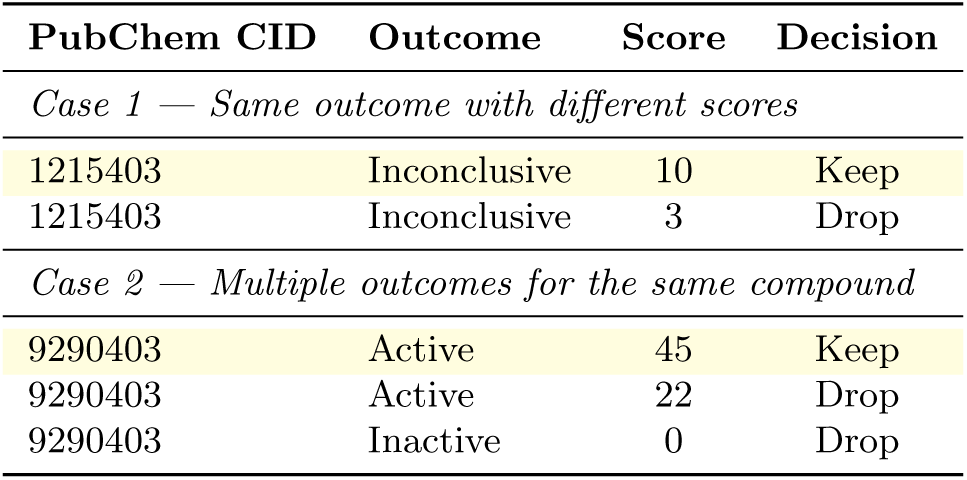
Deduplication examples. Shaded rows are retained; unshaded rows are discarded.

#### Data Integration

When multiple assays from different sources profiled the same cell line, these assay files were merged. For example, the two Caspase-3/7 induction datasets in HepG2 cells, one from Tox21 (AID_1346978) and another from NCATS (AID_654), were combined into a single dataset, after which the same deduplication strategy described above was applied. This procedure yielded our final 18 assays for Caspase-3/7, MMP, and cell viability covering 14 distinct cell lines, as shown in **Table 3**.

**Table 3:**
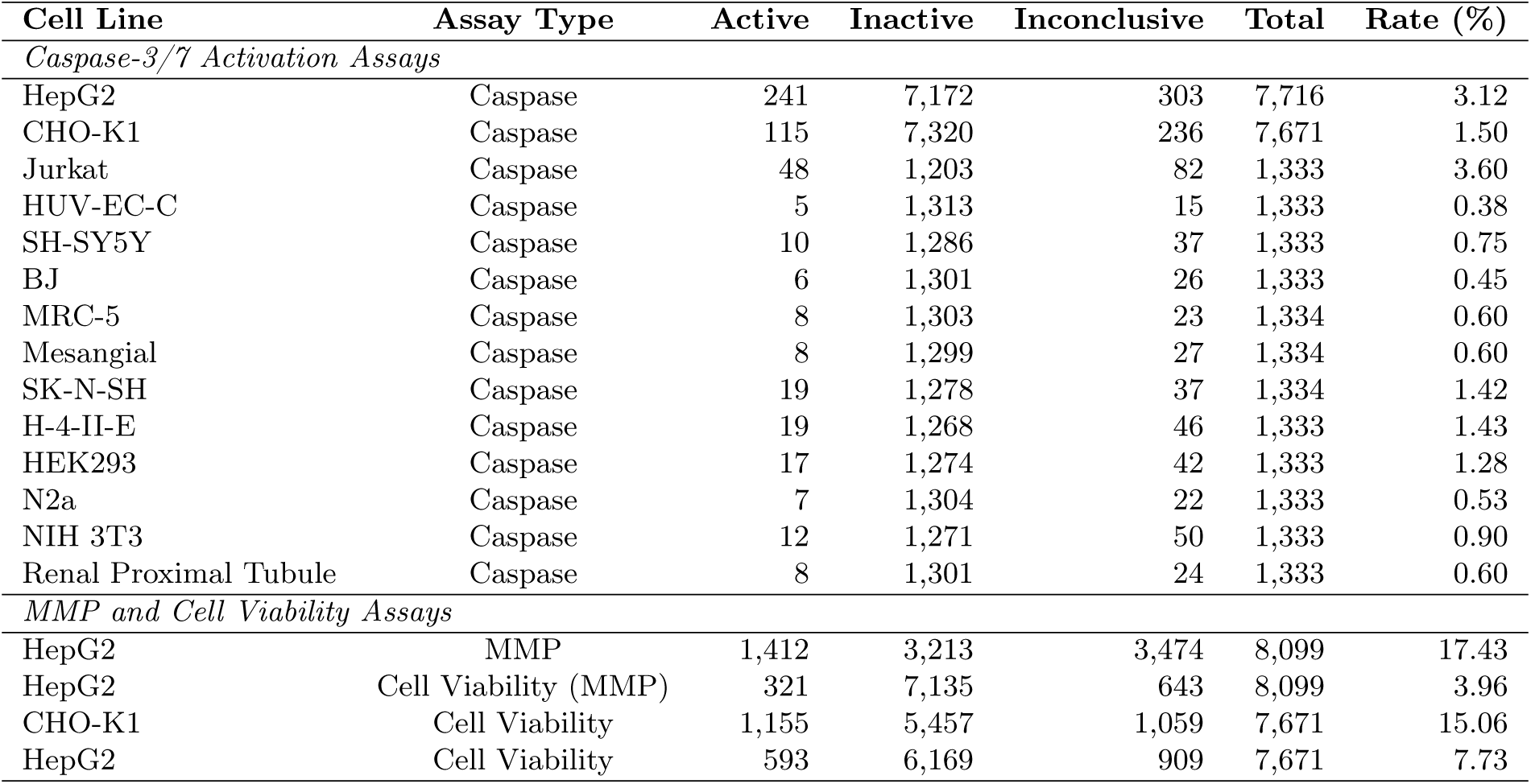
Summary of all 18 qHTS assays: active, inactive, and inconclusive compound counts with activation rates.

As shown in Table 3, Caspase-3/7 activation rates varied substantially across multiple cell lines, ranging from 0.38% in HUV-EC-C to 3.6% in Jurkat cells among the 1,333-compound panels, and reaching 3.12% in the combined HepG2 caspase screen of 7,716 compounds. Mitochondrial and cell viability assays exhibited significantly higher activation rates: the HepG2 mitochondrial membrane assay recorded the highest activation rate at 17.43%, accompanied by a significant inconclusive fraction (42.9% of 8,099 tested).

### Feature Representations and Machine Learning Models

To illustrate the QSAR modeling pipeline, we chose two Caspase-3/7 datasets, HepG2 Caspase-3/7 activation and CHO-K1 Caspase-3/7 activation, to enable a cross-species comparison of Caspase-3/7 induction mechanisms between human hepatic and non-human ovarian cell lines. We also included the MMP dataset to confirm the feasibility of simultaneous monitoring of Caspase-3/7 activation and MMP disruption for novel compounds.

For each dataset, SMILES strings were parsed and converted to canonical form using RDKit [21]; entries that could not be parsed were discarded. Compounds labelled as inconclusive (activity scores 1–39) were excluded during model training. After the processing, the HepG2 Caspase-3/7 dataset retained 7,420 compounds with 249 active (3.4%) and 7,171 inactive (96.6%) compounds, indicating a severe imbalance of 29:1. The CHO-K1 Caspase-3/7 dataset retained 7,435 compounds with 115 active (1.5%) and 7,320 inactive (98.5%) compounds, reflecting a more severe imbalance of 64:1. The mitochondrial toxicity dataset retained 4,625 compounds with 1,412 active (30.5%) and 3,213 inactive (69.5%) compounds.

#### Molecular Feature Representation

We apply a two-track feature engineering strategy to construct both feature representations, molecular fingerprints and molecular graphs, for classical machine learning and graph-based deep learning approaches.

##### Morgan Fingerprints (ECFP4)

Extended-connectivity fingerprints with radius 2 and 2,048 bits (ECFP4) [14] were generated for each compound using RDKit. These circular fingerprints encode the local atomic environment up to two bond distances from each atom and serve as widely used molecular descriptors in QSAR modeling.

##### Molecular Graph Representation

For the graph-based learning track, each compound was encoded as a directed molecular graph with atoms as nodes and bonds as edges; all bonds were represented bidirectionally to enable symmetric message passing. Each atom node was described by a 27-dimensional feature vector encoding atom types (C, N, O, S, F, P, Cl, Br, I, or other; one-hot encoding), degree (0–5; one-hot encoding), formal charge, hybridisation state (sp, sp^2^, sp^3^, or other; one-hot encoding), aromaticity, ring membership, total hydrogen count, and scaled atomic mass. Each bond edge was described by a 6-dimensional feature vector encoding bond type (one-hot encoding), conjugation status, and ring membership.

#### Data Partitioning and Class Balancing

Datasets were partitioned using two sequential stratified splits (random seed 42). First, 20% of each dataset was held out as an external test set. The remaining 80% was then further divided into train and validation subsets by 80/20 split, yielding effective proportions of 64% train, 16% validation, and 20% external test. Class imbalance was addressed by applying random majority-class undersampling exclusively to the training set, producing a 1:1 balanced training set. Validation and test sets were retained at their natural class ratios to reflect realistic deployment conditions. This yielded a balanced HepG2 Caspase-3/7 training set of 318 compounds (159 active, 159 inactive), a balanced CHO-K1 Caspase-3/7 training set of 230 compounds (115 active, 115 inactive), and a balanced mitochondrial training set of 1,808 compounds (904 active, 904 inactive). An alternative undersampling ratio of 5:1 was also evaluated for the Caspase-3/7 datasets and yielded inferior performance; details are provided in Supplementary S1.

#### Classical Machine Learning

Three classical machine learning algorithms were trained and evaluated on the datasets under a grid-search hyperparameter optimisation protocol: Random Forest [22], Support Vector Machine (SVM) with an RBF kernel [23], and XGBoost [24]. All models were trained on the balanced training sets, and hyperparameters were optimised using ROC-AUC on the imbalanced validation set. Full hyperparameter configurations are provided in Supplementary S1.

#### Graph Neural Networks

Four representative message-passing graph neural network architectures were trained and evaluated: Graph Convolutional Network (GCN) [25], Graph Attention Network (GAT) [26], Graph Isomorphism Network (GIN) [27], and GraphSAGE [28]. All four architectures were implemented within a shared modular framework built on PyTorch Geometric [29], using the GCNConv, GATConv, GINConv, and SAGEConv layer primitives, respectively. Input atom features were projected to a hidden dimension via a linear layer with batch normalisation and ReLU activation, followed by three to four message-passing layers, each equipped with batch normalisation, ReLU activation, dropout, and a residual skip connection. Global mean pooling produced a fixed-length graph-level embedding passed to a two-layer MLP classification head trained with binary cross-entropy loss. GAT incorporated multi-head attention with 4 or 8 heads; GIN employed a learnable parameter *ε* and an internal two-layer MLP for neighbour aggregation.

#### Graph Transformers

Graph transformers overcome the locality bias of message-passing GNNs by incorporating global attention that allows every node to attend to every other node simultaneously, avoiding the over-smoothing and limited receptive-field problems inherent in the iterative neighbourhood aggregation [17]. Two representative architectures were implemented: Graphormer [30] and GraphGPS [31]. See their architectural details in Supplementary S1.

Both models share a two-layer MLP classification head (256 *→* 128 *→* 1) trained with class-ratio-weighted binary cross-entropy loss, the Adam optimiser, and a ReduceLROnPlateau scheduler (factor = 0.5, patience = 5). Early stopping (patience = 15, maximum 100 epochs) and a batch size of 64 were used consistently. Full hyperparameter settings are reported in Supplementary S1.

#### Consensus Models

A consensus ensemble strategy was implemented, which aggregates the predictions of individual models to reach a “consensus”, and usually enhances accuracy by mitigating individual model errors. The full consensus prediction was calculated as the arithmetic mean of the predicted active probabilities from all nine trained models: Random Forest, SVM, XGBoost, GCN, GAT, GIN, GraphSAGE, Graphormer, and GraphGPS. Three consensus ensemble variants were evaluated: a classical consensus (RF, SVM, and XGBoost), a GNN consensus (GCN, GAT, GIN, and GraphSAGE), and a GT consensus (Graphormer and GraphGPS).

#### In vivo FDA-approved DILI Prediction

We evaluated our graph-based pipeline on the FDA-approved Drug-Induced Liver Injury (DILI) dataset, comprising 1,111 compounds from DILIst [32] and DILIrank [33] that have been classified as inducing DILI or not and were developed from FDA-approved drug labels. As in Seal et al. [19], the data was partitioned into the *DILI training* set of 888 compounds and the *DILI held-out test* set of 223 compounds using a scaffold-based method (based on the Butina clustering of a bulk Tanimoto fingerprint matrix split with a cutoff threshold of 0.70). This ensures that the DILI test set included a wide variety of compounds that are structurally less similar to those used in training, representing a more challenging split compared to random splits yet matching practical scenarios. We performed SMILES canonicalization using RDKit, removed those compounds that cannot be parsed, and resulted in the DILI training set of 874 compounds (with 553 hepatotoxic and 321 non-hepatotoxic compounds) and the DILI held-out test set of 222 compounds (with 155 hepatotoxic and 67 non-hepatotoxic compounds). We also incorporated the same training data from [19], which uses different in vivo and in vitro data sources, such as liver injury end points for human, preclinical, and animal hepatotoxicity, DILI datasets compiled by various studies, and in vitro assays on mitochondrial toxicity, bile salt export pump inhibition, and the formation of reactive metabolites. This training data is called the *Proxy-DILI* data: after deduplication and removing all 1,111 gold-standard compounds, there are 12,900 unique compounds (active: 4,217; inactive: 8,683).

We selected representative classical models, RF and XGBoost, and graph transformers, Graphormer and GraphGPS, and trained them under two settings: *Proxy-only* with only the *Proxy-DILI* data of 12,900 compounds as the training data and *Proxy+DILI* combining the Proxy-DILI and the DILI training data. Graph transformer models under Proxy-only were pretrained on 12,900 compounds from the Proxy-DILI data using an 80/10/10 split; under Proxy+DILI the pretrained models were further fine-tuning on the DILI training data. Evaluations on the same held-out DILI test set under the same two settings enable a fair comparison of the selected models for their predictive performance for in vivo FDA data. Consensus models (Classic, GTs, and full) were also built and evaluated to confirm their advantages.

#### Applicability Domain

The applicability domain (*AD*) defines the region of chemical space in which a model’s predictions can be considered reliable [18]. An AD was defined independently for each dataset using a 5-nearest-neighbour (5-NN) Jaccard distance criterion applied to the binary ECFP4 training fingerprints. For each training compound *i*, the mean 5-NN Jaccard distance *d̅_i_* was calculated by: 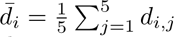 The AD threshold *θ* was set at the mean plus 1.5 standard deviations of the training distribution:

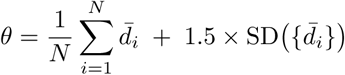

Test compounds with *d̅_q_* = *θ* were flagged as outside the AD and their predictions treated with caution [18].

##### Evaluation Metrics

All models were evaluated at two decision thresholds: a standard threshold of 0.50 (CPT-1) and a high-confidence threshold of 0.80 (CPT-2). A comprehensive set of performance metrics was chosen and reported: *Sensitivity*, *Specificity*, *Correct Classification Rate (CCR* = *(sensitivity* + *specificity) /*2), *Receiver Operating Characteristic - Area Under Curve (ROC-AUC)*, *Area Under the Precision-Recall Curve (PR-AUC)*, *Matthews correlation coefficient (MCC)*, and *F1-score*. We choose the widely adopted ROC-AUC and F1-score as the primary performance measures.

## Results and discussions

### Prediction Results of QSAR Models

#### Mitochondrial Toxicity Dataset

Mitochondrial membrane potential (MMP) plays a central role in maintaining cellular energy homeostasis and survival, and its loss is an early initiating event in the intrinsic apoptotic pathway leading to Caspase-3/7 activation.

**Figure 2**(B) summarizes the F1 and ROC-AUC scores across all tested models. **Table 4** presents the full test set performance of all thirteen models at both decision thresholds for the mitochondrial toxicity dataset. **Figure 3** shows their corresponding AUC curves. Remarkably, *the full consensus model achieved the highest ROC-AUC of 0.92 and the best F1 of 0.77* at CPT-1. We observed that all GNN architectures outperformed classical models across both thresholds, due to their powerful ability to capture structural patterns. Additionally, all the consensus ensembles surpassed their respective constituent models, showing the advantages of combining multiple approaches to reduce variance and bias. While individual graph transformers have suboptimal performance in ROC-AUC and F1, their predictions tend to well complement each other and their consensus model (consensus-GT) achieved significant performance boost of 0.025 in ROC-AUC and 0.023 in F1, making it the second best model in our test while being less computationally expensive than the full consensus.

**Figure 2:**
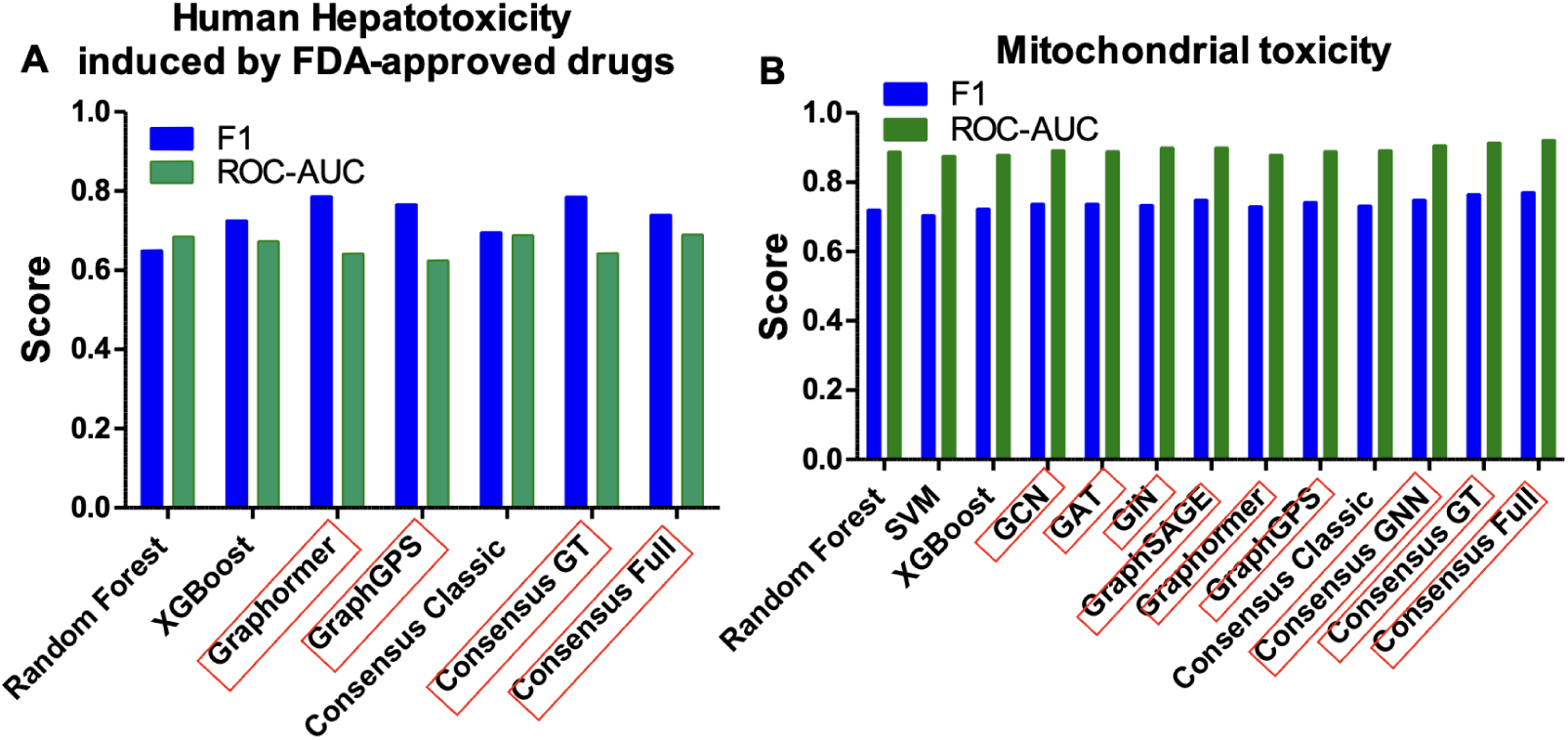
(A) F1 and ROC-AUC scores for all tested models under the Proxy+DILI setting on the FDA-approved DILI held-out test set. (B) F1 and ROC-AUC scores for all tested models on the mitochondrial toxicity dataset. Graph-based models are highlighted.

**Figure 3:**
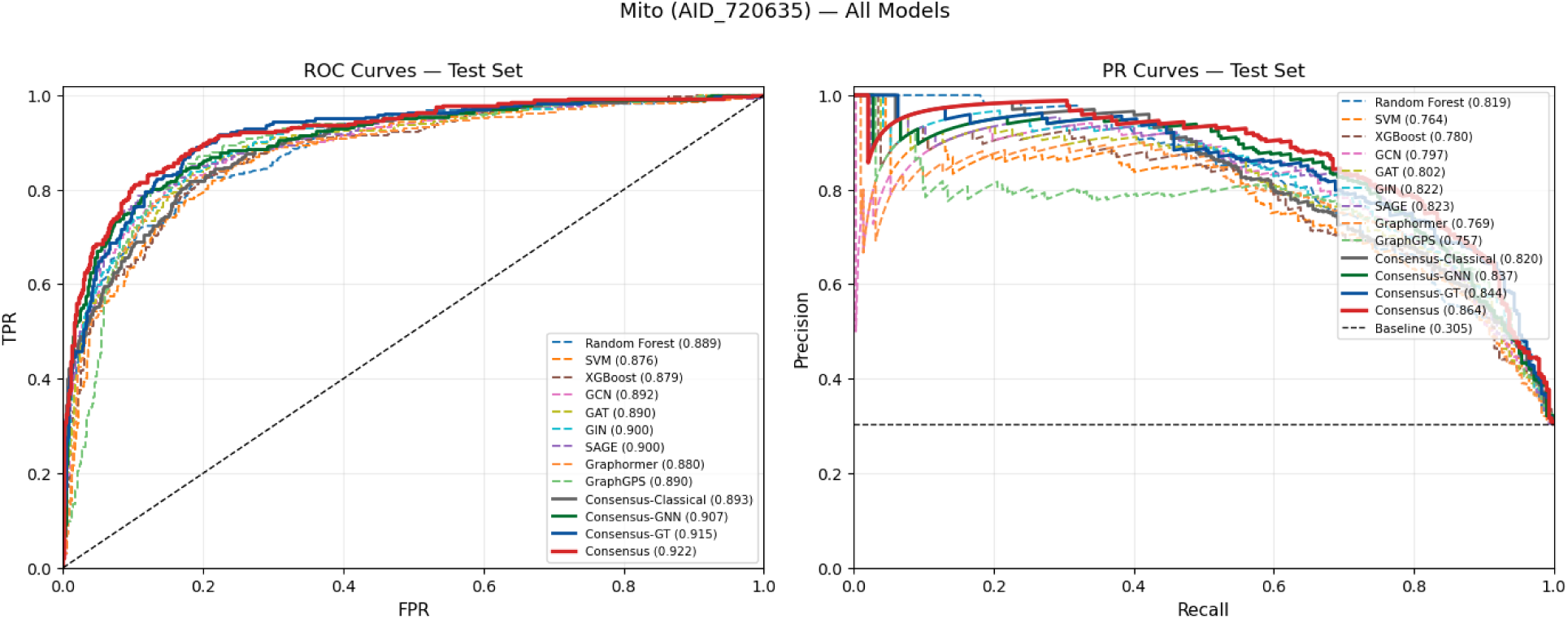
ROC-AUC and PR-AUC curves for all thirteen models on the mitochondrial toxicity dataset. AUC values are shown in the legend.

**Table 4:** Performance of various QSAR models in predicting mitochondrial toxicity — Test Set Performance (64/16/20 stratified split, 1:1 downsampled train)

| Model | CPT | Sensitivity (%) | Specificity (%) | CCR (%) | ROC-AUC | PR-AUC | MCC | Bal-Acc (%) | F1 |
| --- | --- | --- | --- | --- | --- | --- | --- | --- | --- |
| *Classical Machine Learning* |  |  |  |  |  |  |  |  |  |
| Random Forest | 0.5 | 81.9 | 80.2 | 81.1 | 0.8893 | 0.8192 | 0.5875 | 81.1 | 0.7219 |
|  | 0.8 | 42.6 | **98.1** | 70.3 | 0.8893 | 0.8192 | 0.5355 | 70.3 | 0.5797 |
| SVM | 0.5 | 79.4 | 79.9 | 79.7 | 0.8763 | 0.7643 | 0.5626 | 79.7 | 0.7055 |
|  | 0.8 | 55.3 | 94.4 | 74.9 | 0.8763 | 0.7643 | 0.5644 | 74.9 | 0.6582 |
| XGBoost | 0.5 | 81.2 | 81.2 | 81.2 | 0.8795 | 0.7805 | 0.5922 | 81.2 | 0.7247 |
|  | 0.8 | 61.7 | 91.8 | 76.7 | 0.8795 | 0.7805 | 0.5719 | 76.7 | 0.6837 |
| *Graph Neural Networks* |  |  |  |  |  |  |  |  |  |
| GCN | 0.5 | 82.6 | 82.0 | 82.3 | 0.8924 | 0.7972 | 0.6134 | 82.3 | 0.7385 |
|  | 0.8 | 66.3 | 93.8 | 80.0 | 0.8924 | 0.7972 | 0.6428 | 80.0 | 0.7348 |
| GAT | 0.5 | 83.7 | 81.2 | 82.4 | 0.8899 | 0.8019 | 0.6134 | 82.4 | 0.7387 |
|  | 0.8 | 54.6 | 96.4 | 75.5 | 0.8899 | 0.8019 | 0.5972 | 75.5 | 0.6710 |
| GIN | 0.5 | 84.4 | 80.1 | 82.2 | 0.9005 | 0.8223 | 0.6071 | 82.2 | 0.7346 |
|  | 0.8 | 69.1 | 91.9 | 80.5 | 0.9005 | 0.8223 | 0.6354 | 80.5 | 0.7372 |
| SAGE | 0.5 | 84.0 | 82.4 | 83.2 | 0.9001 | 0.8233 | 0.6309 | 83.2 | 0.7500 |
|  | 0.8 | 73.8 | 92.1 | 82.9 | 0.9001 | 0.8233 | **0.6749** | 82.9 | 0.7689 |
| *Graph Transformers* |  |  |  |  |  |  |  |  |  |
| Graphormer | 0.5 | 79.4 | 83.4 | 81.4 | 0.8798 | 0.7687 | 0.6030 | 81.4 | 0.7308 |
|  | 0.8 | 57.4 | 93.9 | 75.7 | 0.8798 | 0.7687 | 0.5735 | 75.7 | 0.6708 |
| GraphGPS | 0.5 | 87.2 | 79.2 | 83.2 | 0.8902 | 0.7569 | 0.6213 | 83.2 | 0.7432 |
|  | 0.8 | **79.1** | 87.6 | **83.3** | 0.8902 | 0.7569 | 0.6536 | **83.3** | 0.7624 |
| *Consensus Ensembles* |  |  |  |  |  |  |  |  |  |
| Consensus-Classical | 0.5 | 81.9 | 81.8 | 81.9 | 0.8932 | 0.8197 | 0.6055 | 81.9 | 0.7333 |
|  | 0.8 | 52.5 | 96.3 | 74.4 | 0.8932 | 0.8197 | 0.5768 | 74.4 | 0.6520 |
| Consensus-GNN | 0.5 | 84.4 | 82.1 | 83.3 | 0.9073 | 0.8366 | 0.6303 | 83.3 | 0.7496 |
|  | 0.8 | 65.6 | 95.3 | 80.5 | 0.9073 | 0.8366 | 0.6641 | 80.5 | 0.7445 |
| Consensus-GT | 0.5 | **87.6** | 82.0 | 84.8 | 0.9152 | 0.8439 | 0.6557 | 84.8 | 0.7659 |
|  | 0.8 | 63.8 | 95.0 | 79.4 | 0.9152 | 0.8439 | 0.6446 | 79.4 | 0.7287 |
| Consensus (Full) | 0.5 | 86.5 | **83.5** | **85.0** | **0.9225** | **0.8636** | **0.6648** | **85.0** | **0.7722** |
|  | 0.8 | 57.8 | 97.5 | 77.7 | **0.9225** | **0.8636** | 0.6446 | 77.7 | 0.7072 |

Our results show that when the number of known toxic compounds is relatively large, such as over 1,400 in the Tox21 mitochondrial toxicity assay, it is promising to build a QSAR model with strong performance in predicting mitochondrial toxicity. Validation on different cell lines and organs/tissues beyond hepatotoxicity is needed as the relevant data is available.

#### HepG2 and CHO-K1 Caspase-3/7 Datasets

The HepG2 Caspase-3/7 dataset poses a substantially harder prediction problem than the mitochondrial endpoint, because of its small number of toxic compounds and a severe nontoxic/toxic class imbalance of 28.8:1. Complete per-model results at both decision thresholds are provided in Supplementary Table S1. Across all models, ROC-AUC values ranged from 0.69 to 0.8 and F1 ranged from 0.08 to 0.18, compared with 0.88–0.92 and 0.58–0.77 on the mitochondrial dataset, respectively. The significant reduction on F1 scores reflects the true challenge under class imbalance. Consistent with the mitochondrial data, *the full consensus achieved the second highest ROC-AUC of 0.79 and the highest F1 of 0.16* (at CPT1). Interestingly, Random Forest attained the highest ROC-AUC of 0.8 and an F1 of 0.15, demonstrating a comparable performance compared to the full consensus under the class imbalance scenario. Graphormer recorded a low performance in this data for some reason; however, the full consensus model is robust to the abnormality, maintaining its status as a top-ranked approach.

The CHO-K1 Caspase-3/7 dataset is an even more challenging endpoint, with only 115 toxic compounds and a class imbalance of 63.7:1 in the curated dataset. Complete per-model results are provided in Supplementary Table S2. ROC-AUC and F1 values are similarly low with a large variance, ranging from 0.67 to 0.83 and from 0.04 to 0.21 respectively. The full consensus achieved a ROC-AUC of 0.76, and an F1 of 0.13 (at CPT-2). Remarkably, graph transformers performed the best among individual methods in this setting. Their consensus model consensus-GT further boosts the ROC-AUC to 0.83 and the F1 to 0.18, making it the best model in this data. SVM showed a good performance with ROC-AUC of 0.78 and the highest F1 of 0.21.

Our results show that when the number of known toxic compounds is small and the class imbalance is severe, such as in these Caspase-3/7 datasets, it is challenging to build a QSAR model with strong prediction performance, especially on reducing false negatives. The full consensus and consensus-GT usually performed relatively the best while classic models Random Forest and SVM also showed relatively good performance.

##### Impacts of Applicability Domain

The applicability domain threshold was set at 0.65. Out of the 925 test compounds, 514 (55.6%) were classified as within the applicability domain and 411 (44.4%) as outside (Figure 4, left). Table 5 compares the consensus model performance on the full test set and the AD-filtered subset in the mitochondrial toxicity dataset. Restricting predictions to within-domain compounds consistently improved all metrics: sensitivity increased from 86% to 90.9%, CCR from 85% to 87.9%, ROC-AUC from 0.91 to 0.93, and MCC from 0.67 to 0.74 (Figure 4, right). Specificity remained stable (84.0% vs. 84.8%), indicating that the performance gain was driven primarily by improved identification of active compounds. Compounds outside the domain represent structurally novel scaffolds that warrant experimental confirmation before relying on model predictions.

**Figure 4:**
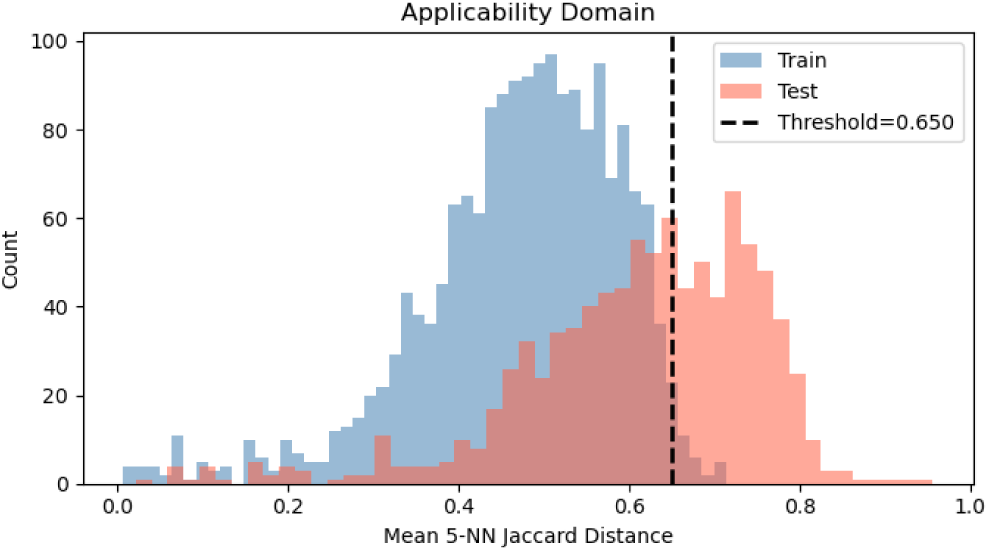
Applicability domain analysis: proportion of the 925 test compounds within (55.6%) and outside (44.4%) the AD.

**Table 5:** Consensus model performance on the full test set versus the AD-filtered subset (mitochondrial toxicity dataset).

| Metric | Full Test Set | AD-Filtered |
| --- | --- | --- |
| Sensitivity | 86% | 90.9% |
| Specificity | 84% | 84.8% |
| CCR | 85% | 87.9% |
| ROC-AUC | 0.91 | 0.93 |
| MCC | 0.67 | 0.74 |

#### In vivo FDA-approved DILI Dataset & Structural Alerts from Graph Transformers

**Figure 2**(A) presents the F1 and ROC-AUC scores under the Proxy+DILI setting across all tested models. **Table 6** reports experimental results for all models in the held-out test set of 222 compounds from the FDA-approved DILI benchmark. **Figure 5** shows their ROC-AUC and PR-AUC curves. The inclusion of DILI training data consistently improved the prediction performance of all tested models compared to the Proxy-only setting. This matches with our expectation since the proxy-DILI data do not contain compounds labeled by rigorous FDA study and the DILI training data can provide more accurate supervision on the prediction of DILI test data. In particular, RF achieved a decent ROC-AUC of 0.69 and an F1 of 0.65 at a decision threshold of 0.5 while graphormer recorded the highest F1 of 0.79 and an ROC-AUC of 0.64. The full consensus model further improved the ROC-AUC to 0.69 while maintaining a decent F1 of 0.74.

**Figure 5:**
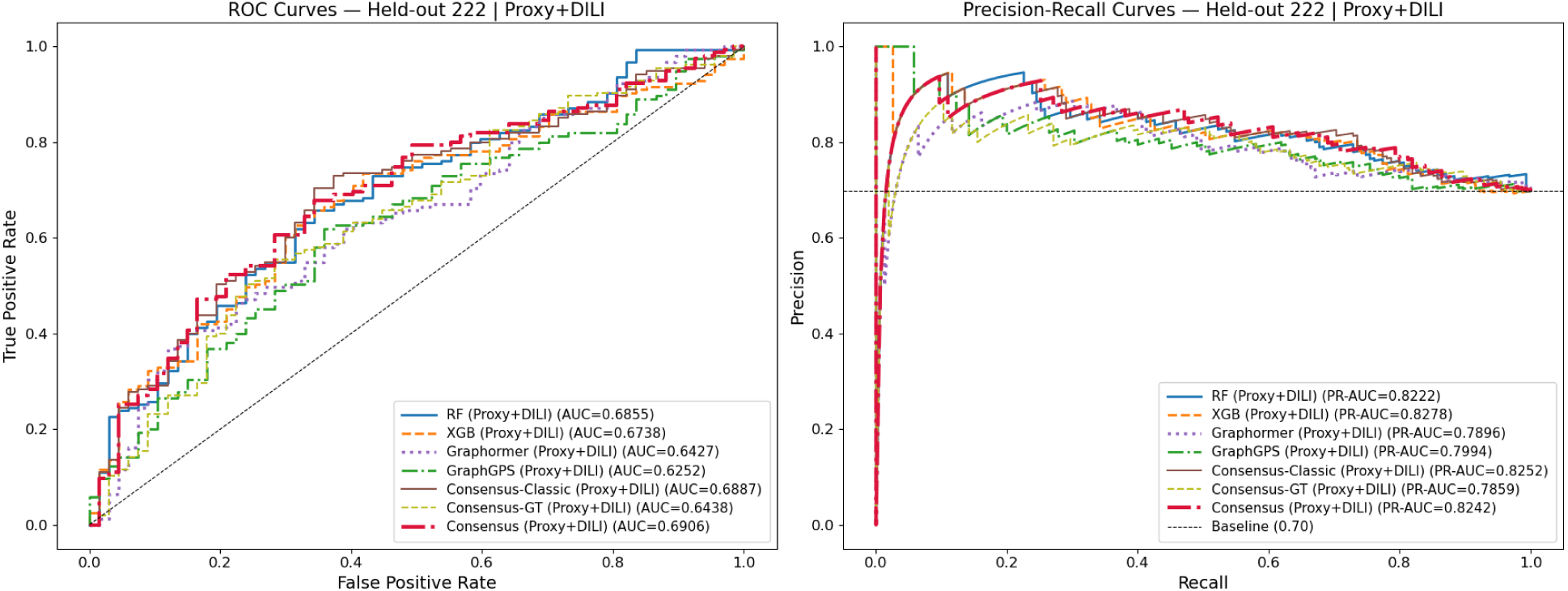
ROC-AUC and PR-AUC curves for all models under the Proxy+DILI setting on the FDA DILI held-out test set

**Table 6:** Performance of all models on the FDA-approved DILI held-out test set at decision threshold of 0.5 (222 compounds after canonicalization; 155 hepatotoxic, 67 non-hepatotoxic). Sens = sensitivity (%), Spec = specificity (%). Bold indicates best value per column.

| Model | Sens | Spec | CCR | ROC-AUC | PR-AUC | MCC | F1 |
| --- | --- | --- | --- | --- | --- | --- | --- |
| *Classical ML* |  |  |  |  |  |  |  |
| RF (Proxy-only) | 52.3 | **77.6** | 64.9 | 0.6711 | 0.8138 | 0.277 | 0.645 |
| RF (Proxy+DILI) | 53.5 | 74.6 | 64.1 | 0.6855 | 0.8222 | 0.260 | 0.651 |
| XGB (Proxy-only) | 65.2 | 61.2 | 63.2 | 0.6580 | 0.8057 | 0.245 | 0.716 |
| XGB (Proxy+DILI) | 65.8 | 64.2 | **65.0** | 0.6738 | **0.8278** | **0.278** | 0.726 |
| *Graph Transformers* |  |  |  |  |  |  |  |
| Graphormer (Proxy-only) | 74.8 | 38.8 | 56.8 | 0.6507 | 0.7924 | 0.138 | 0.744 |
| Graphormer (Proxy+DILI) | **85.8** | 25.4 | 55.6 | 0.6427 | 0.7896 | 0.135 | **0.787** |
| GraphGPS (Proxy-only) | 79.4 | 28.4 | 53.9 | 0.6001 | 0.7757 | 0.084 | 0.755 |
| GraphGPS (Proxy+DILI) | 81.9 | 26.9 | 54.4 | 0.6252 | 0.7994 | 0.100 | 0.767 |
| *Ensemble* |  |  |  |  |  |  |  |
| Consensus-Classic (Proxy-only) | 58.7 | 67.2 | 62.9 | 0.6697 | 0.8134 | 0.238 | 0.679 |
| Consensus-Classic (Proxy+DILI) | 60.6 | 68.7 | 64.7 | 0.6887 | 0.8252 | 0.269 | 0.696 |
| Consensus-GT (Proxy-only) | 71.0 | 46.3 | 58.6 | 0.6394 | 0.7846 | 0.167 | 0.731 |
| Consensus-GT (Proxy+DILI) | 83.2 | 34.3 | 58.8 | 0.6438 | 0.7859 | 0.194 | 0.787 |
| Consensus (Proxy-only) | 67.1 | 59.7 | 63.4 | 0.6775 | 0.8093 | 0.250 | 0.727 |
| Consensus (Proxy+DILI) | 69.0 | 59.7 | 64.4 | **0.6906** | 0.8242 | 0.270 | 0.741 |

**Figures 6, 7, and 8** present SubgraphX explanations for the top-6 true positives identified by each graph transformer on the FDA-approved DILI held-out test set. SubgraphX [34] identifies the minimal connected subgraph most responsible for the model’s prediction; highlighted atoms (red) represent the structural alert driving the hepatotoxic classification. Graphormer assigned highest confidence to halothane (prob=0.808), while GraphGPS ranked sorafenib highest (prob=0.718). The consensus panel shows the top-6 compounds independently flagged by both models, where cross-model agreement on known hepatotoxins suggests the graph transformers have learned chemically meaningful structural alerts.

**Figure 6:**
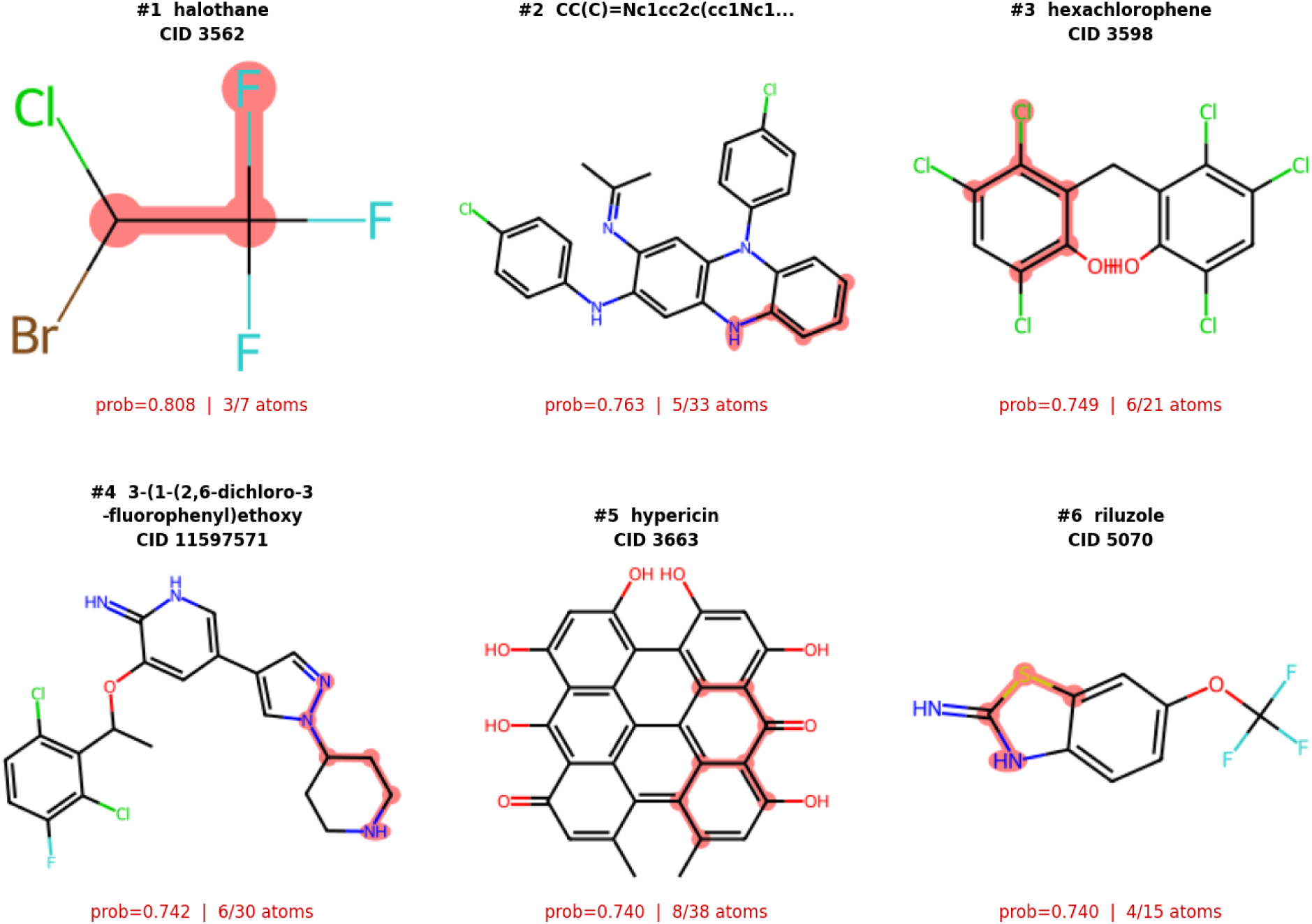
Graphormer - Top 6 DILI true positives from the FDA held-out test set. Red subgraphs indicate the atoms most responsible for the hepatotoxic prediction (SubgraphX). prob = model confidence; atoms = subgraph size.

**Figure 7:**
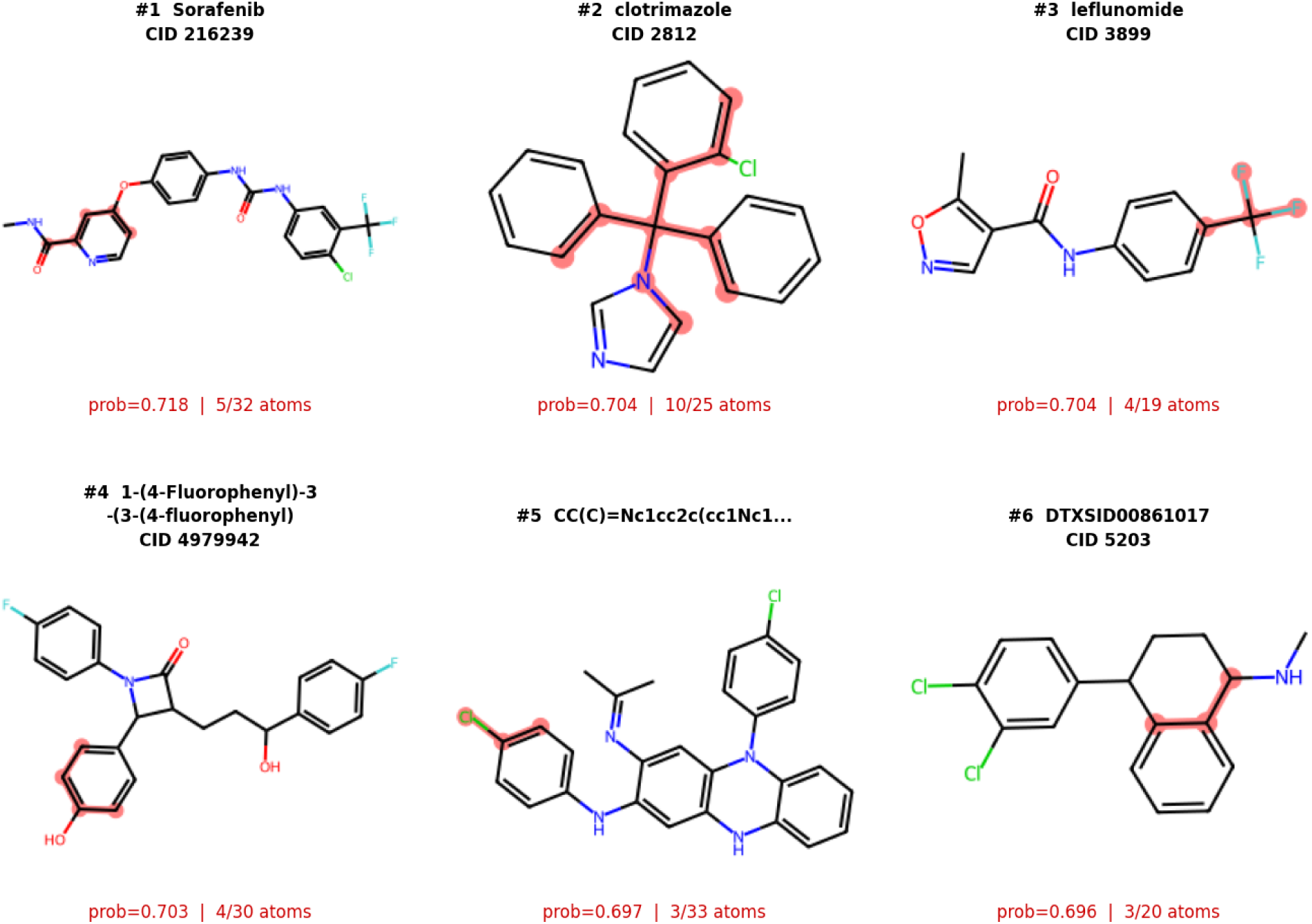
GraphGPS - Top 6 DILI true positives from the FDA held-out test set, with SubgraphX toxic subgraphs highlighted in red.

**Figure 8:**
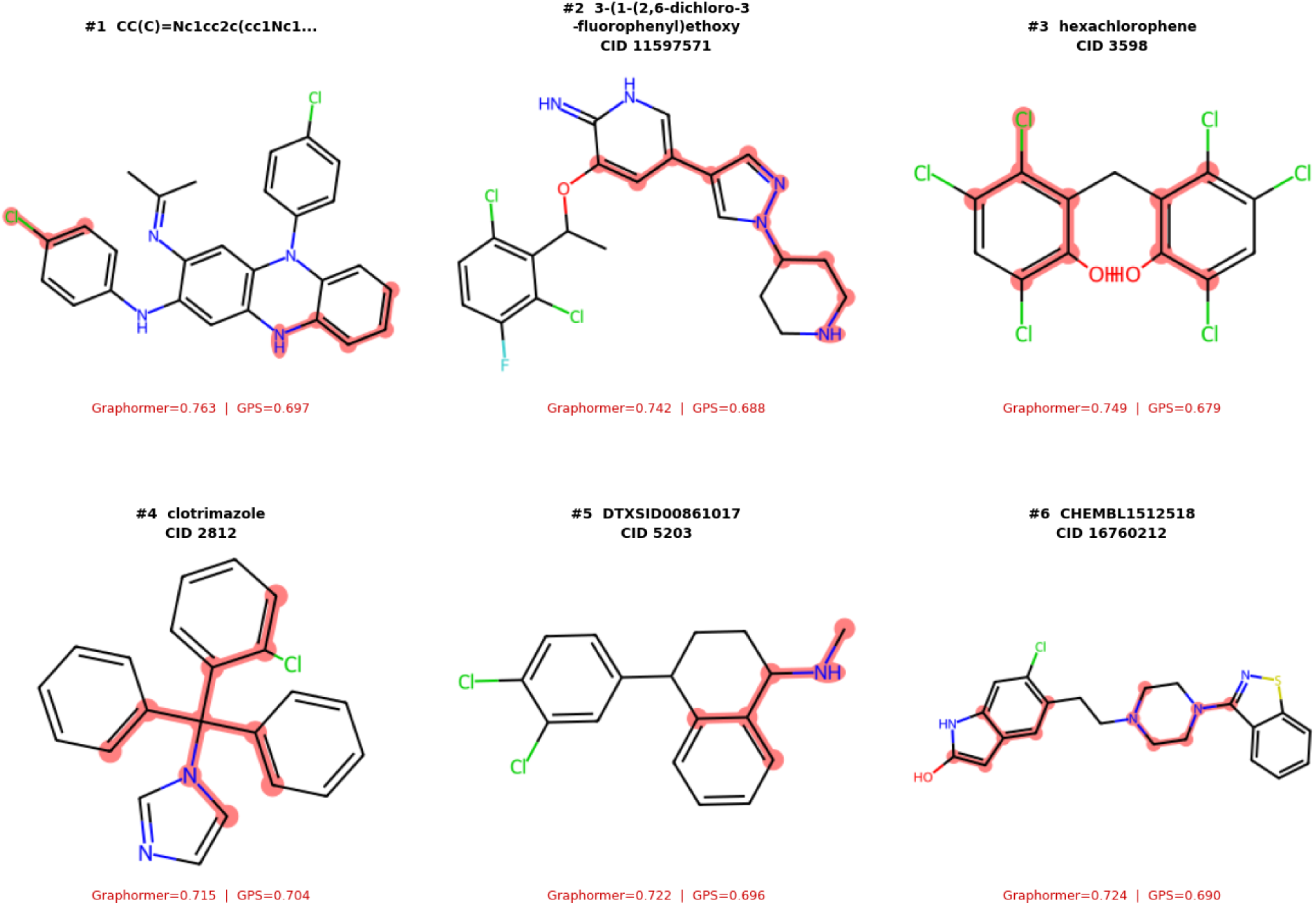
Consensus DILI compounds flagged by both Graphormer and GraphGPS. Red subgraphs highlight the toxic structural motifs; confidence scores from each model are shown below each compound.

Our proposed graph-based QSAR modeling pipeline significantly outperformed the previously best model, the multi-feature DILIPredictor [35] with ROC-AUC of 0.63 and F1 of 0.65 on the *same* held-out test set, which even additionally incorporated physicochemical descriptors and pharmacokinetic parameters. Based on the promising preliminary results, we will further explore how to leverage abundant existing hepatotoxicity studies from PubMed literature and use advanced LLM-driven approaches for improving DILI prediction of novel compounds.

### Cytotoxicity Activation Behavior and Structural Analysis

#### Mitochondrial Membrane Potential (MMP) Disruption Pathway and Caspase-3/7 Activation

Since both Caspase-3/7 activation and mitochondrial activation data are available in HepG2 cells, this enables pairwise comparison between compounds tested in both assays to identify activation patterns and analyze the underlying apoptosis pathway. As shown in **Table 7**, 144 of 3,761 compounds (3.8%) were active in both Caspase and mitochondrial assays, confirming the mitochondrial pathway leading to apoptosis. Seventeen compounds (0.5%) were Caspase-active but mitochondria-inactive. Compared with 3.8% dual-active category, this indicates a majority of the Caspase activation is caused by the mitochondrial activity. It is consistent with the general understanding on the mitochondrial pathway to apoptosis. We now analyze representative substructures presented in “both active” compounds vs “both inactive” compounds in Table 7.

**Table 7:**
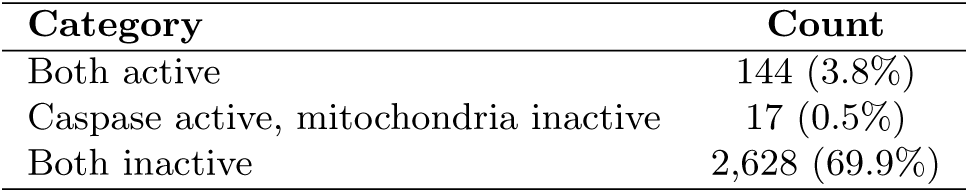
Pairwise comparison of Caspase-3/7 and mitochondrial activity in HepG2 cells.

##### Structural Motifs in Dual-Active Versus Dual-Inactive Compounds

To identify structural features associated with compounds that fully engage the caspase–mitochondrial apoptotic cascade, a *Breaking Retrosynthetically Interesting Chemical Substructures (BRICS)* fragment enrichment analysis was performed contrasting 144 dual-active compounds against 2,628 dual-inactive compounds. Specifically, the BRICS method [36] first decomposes these compounds into chemically meaningful fragments. Then a one-sided Fisher’s exact test with Benjamini–Hochberg correction (*α* = 0.05) is applied to each fragment to decide whether it is significantly presented in dual-active compared to dual-inactive compounds. The one-sided Fisher’s exact test was chosen because fragment occurrence is binary per molecule and expected cell counts are frequently below five in targeted chemical subsets, making the chi-squared approximation unreliable; the one-sided formulation tests the directional hypothesis that a fragment appears more often in actives than inactives. This analysis identified 16 significantly enriched fragments in the dual-active foreground compared to the dual-inactive background. (**Figure 9**; **Table 8**). These fragments can be considered as defining chemical substructures for dual-active compounds and can be organized into four structural families.

The *phenolic* family comprised three fragments and produced the strongest signal. A para-hydroxyphenyl motif was the top-ranked fragment overall, that is the one with smallest adjusted *p*-value (*p*_adj_ = 5.0 *×* 10*^−^*^7^). It is presented in 8.3% of dual-active compounds versus 0.7% of dual-inactive compounds. A disubstituted hydroxyphenyl variant and an *m*-cresol motif in the phenolic family further reinforced this finding. Together, these fragments establish hydroxyl-substituted aromatic rings as the most statistically reliable structural predictor of caspase–mitochondrial co-activation.

**Figure 9:**
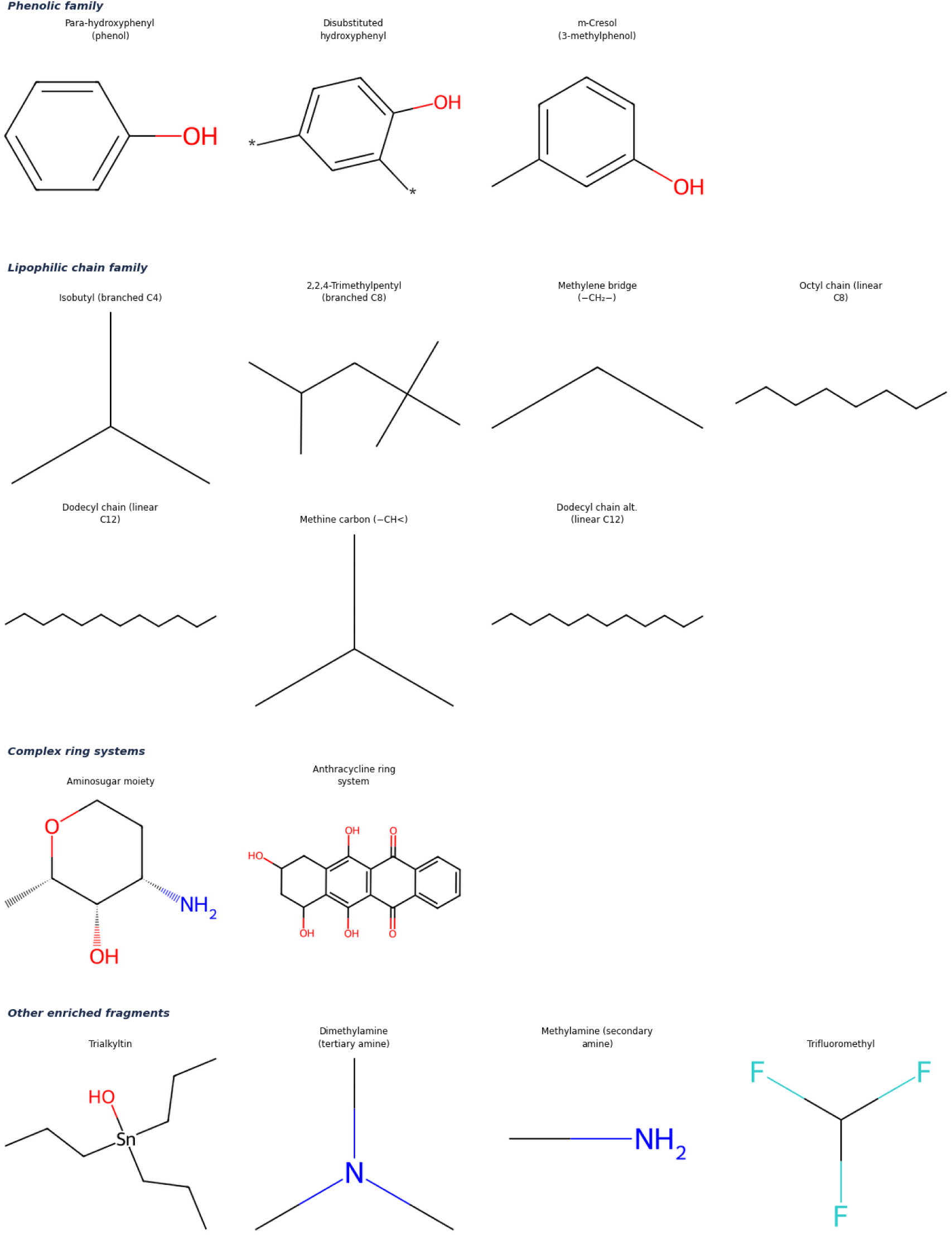
All 16 BRICS fragments significantly enriched in **dual-active** (Caspase + mitochondrial) versus dual-inactive compounds, grouped by the structural family. Full enrichment statistics, including odds ratios, adjusted *p*-values, and per-group prevalence counts, are provided in Table 8.

**Table 8:**
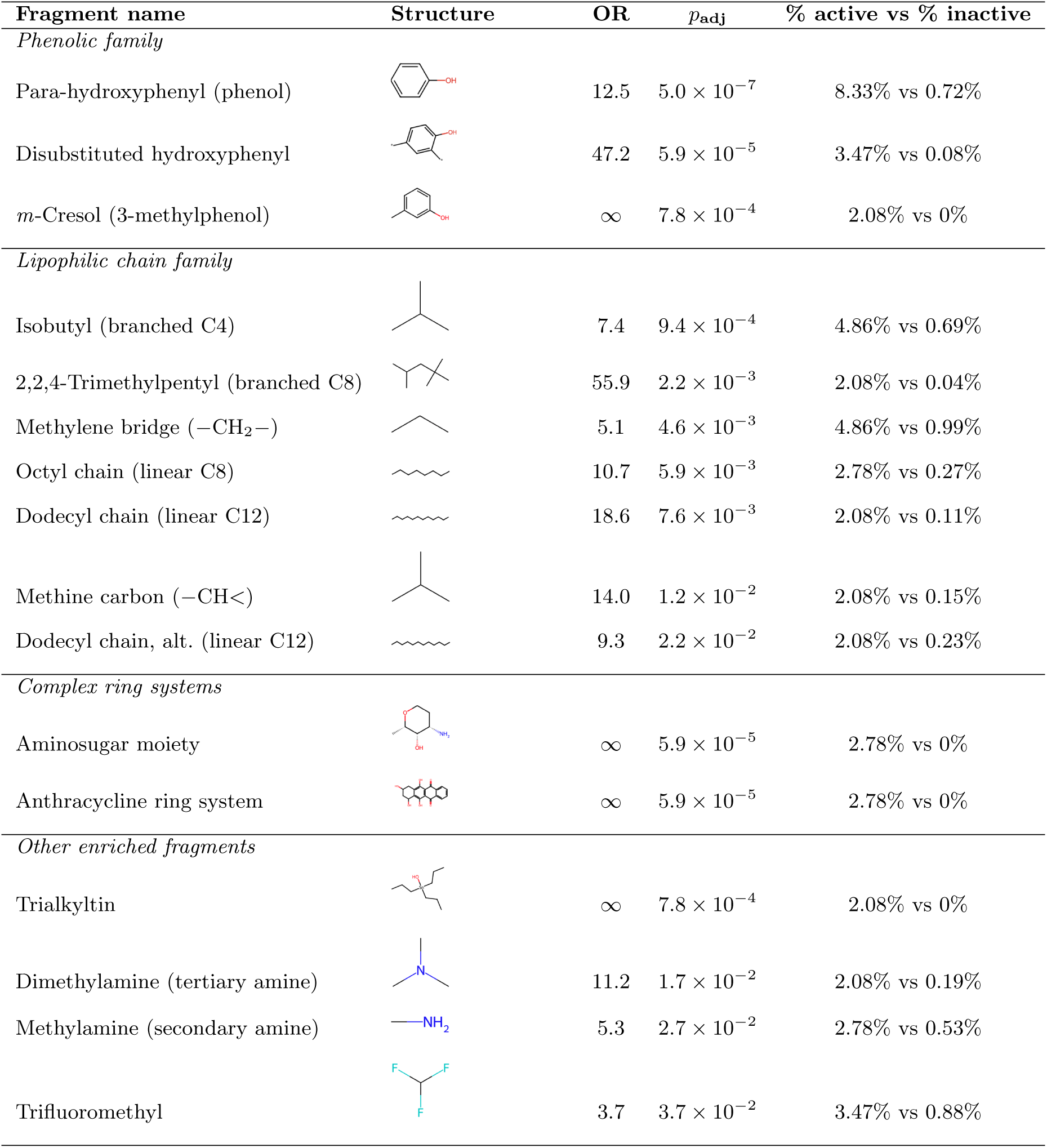
All 16 BRICS fragments significantly enriched in dual-active (Caspase + mitochondrial) versus dual-inactive compounds (*n*_active_ = 144, *n*_inactive_ = 2,628), grouped by structural family and ranked by adjusted *p*-value (Benjamini–Hochberg, *α* = 0.05). OR = *∞* denotes fragments absent from the dual-inactive set. % active and % inactive denote percentages of dual-active and dual-inactive compounds with a fragment respectively.

| Fragment name | Structure | OR | $p_{\text{adj}}$ | % active vs % inactive |
| --- | --- | --- | --- | --- |
| *Phenolic family* |  |  |  |  |
| Para-hydroxyphenyl (phenol) | | 12.5 | $5.0 \times 10^{-7}$ | 8.33% vs 0.72% |
| Disubstituted hydroxyphenyl | | 47.2 | $5.9 \times 10^{-5}$ | 3.47% vs 0.08% |
| *m*-Cresol (3-methylphenol) | | ∞ | $7.8 \times 10^{-4}$ | 2.08% vs 0% |
| *Lipophilic chain family* |  |  |  |  |
| Isobutyl (branched C4) | | 7.4 | $9.4 \times 10^{-4}$ | 4.86% vs 0.69% |
| 2,2,4-Trimethylpentyl (branched C8) | | 55.9 | $2.2 \times 10^{-3}$ | 2.08% vs 0.04% |
| Methylene bridge ($-\text{CH}_2-$) | | 5.1 | $4.6 \times 10^{-3}$ | 4.86% vs 0.99% |
| Octyl chain (linear C8) | | 10.7 | $5.9 \times 10^{-3}$ | 2.78% vs 0.27% |
| Dodecyl chain (linear C12) | | 18.6 | $7.6 \times 10^{-3}$ | 2.08% vs 0.11% |
| Methine carbon ($-\text{CH}<$) | | 14.0 | $1.2 \times 10^{-2}$ | 2.08% vs 0.15% |
| Dodecyl chain, alt. (linear C12) | | 9.3 | $2.2 \times 10^{-2}$ | 2.08% vs 0.23% |
| *Complex ring systems* |  |  |  |  |
| Aminosugar moiety | | ∞ | $5.9 \times 10^{-5}$ | 2.78% vs 0% |
| Anthracycline ring system | | ∞ | $5.9 \times 10^{-5}$ | 2.78% vs 0% |
| *Other enriched fragments* |  |  |  |  |
| Trialkyltin | | ∞ | $7.8 \times 10^{-4}$ | 2.08% vs 0% |
| Dimethylamine (tertiary amine) | | 11.2 | $1.7 \times 10^{-2}$ | 2.08% vs 0.19% |
| Methylamine (secondary amine) | | 5.3 | $2.7 \times 10^{-2}$ | 2.78% vs 0.53% |
| Trifluoromethyl | | 3.7 | $3.7 \times 10^{-2}$ | 3.47% vs 0.88% |

The *lipophilic chain* family comprised seven fragments, including branched and linear alkyl chains from C4 to C12. The branched C8 chain (2,2,4-trimethylpentyl) showed the strongest enrichment within this group, followed by linear C12 and octyl. A methylene bridge and a methine carbon were also enriched, reflecting the branched connectivity typical of this compound class. The co-occurrence of phenolic and lipophilic families in the same foreground indicates that dual apoptotic activity is accessible via at least two structurally independent routes.

Two complex *ring system* fragments were identified exclusively in dual-active compounds (and not presented in dual-inactive compounds): an aminosugar moiety and an anthracycline-derived ring system (both with *p*_adj_ = 5.9 *×* 10*^−^*^5^). The remaining four enriched fragments spanned distinct chemical classes: a *trialkyltin group*, a *tertiary dimethylamine*, a *secondary methylamine*, and a *trifluoromethyl group*. It should be notable that the reverse analysis—fragments enriched in dual-inactive compounds compared to dual-active compounds—returned no significant hits, indicating that structural inactivity has no defining chemical motif.

Compounds featuring lipophilic alkyl chains and cationic groups, such as delocalized lipophilic cations, were reported to selectively accumulate within mitochondria due to the negative potential characteristic of the mitochondrial inner membrane; here, they can directly interfere with mitochondrial function [37]. Notably, enhancing lipophilicity through the introduction of longer alkyl chains increases a compound’s affinity for membranes as well as the extent of its accumulation within mitochondria. This heightened lipophilicity correlates positively with the compound’s capacity to reduce the mitochondrial inner membrane potential (Δ*ψ_m_*); the underlying mechanism involves disrupting the integrity of the inner mitochondrial membrane and increasing proton leakage [38]. These lipophilic cationic compounds can intercalate into mitochondrial membranes, disrupt the function of respiratory chain complexes, and ultimately trigger the collapse of Δ*ψ_m_* [38], leading to the activation of caspases-3 and -7. Furthermore, phenolic groups and redox-active moieties can also induce mitochondrial dysfunction and thereby further destabilize the membrane potential by promoting oxidative stress and mitochondrial damage [39]. This study reveals that phenolic compounds bearing a para-hydroxyphenyl motif, as well as members of the lipophilic chain family specifically those possessing structural amphiphilicity (combining hydrophobic and polar characteristics) or ionophore-like structures can disrupt the ion gradients across the mitochondrial membrane. This disruption directly dissipates Δ*ψ_m_* and triggers mitochondrial swelling and dysfunction. In contrast, compounds lacking sufficient lipophilicity or specific mitochondrial-targeting features exhibit lower levels of mitochondrial accumulation and, consequently, exert a relatively weaker disruptive effect on Δ*ψ_m_*. Moreover, amine-containing groups, such as dimethylamine, when protonatable or integrated into an ionizable molecular scaffold, can influence a compound’s mitochondrial-targeting behavior by modulating the balance between membrane permeability and charge-driven accumulation. Relevant experimental studies utilizing lipophilic cations coupled with ionizable groups (including amino groups) have demonstrated that the efficiency of compound uptake within mitochondria depends on the compound’s *pK_a_* value, its protonation state, and the pH gradient across the mitochondrial membrane; this directly confirms that amino functional groups play a pivotal role in determining mitochondrial targeting behavior [40]. This study suggests that, specifically for dual-activity compounds, namely those comprising an aminosugar moiety and an anthracycline-derived ring likely key physicochemical determinants may be responsible for their ability to target mitochondria.

Our results mainly support findings of the previous study on structural features for Caspase activation by Huang *et al.* [12], which showed that cyclic alkyl ketones and alkyl halides were the most dominant features in the caspase-activating compounds. Our results further show a collection of structural motifs, such as lipophilic alkyl chains, phenolic family, and complex ring systems, may be responsible for dual-mitochontrial-caspase activation.

#### Caspase Activations and Cell-Line-Specific Structural Motifs

The only study on Caspase-3/7 activation prior to the present work is Huang et al. [12], who considered “pan-Caspase-3/7 activator” by combining all 13 different cell types and taking their average, and also identified structural features for the pan-caspase toxicity. They significantly ignored the impacts of diverse cell types on the Caspase-3/7 activation as cells of different types can exhibit completely different responses to the same event. Hence, they cannot offer finer-level cross-cell-line analysis and identify the activation differences among the tested cell lines. The weighted feature significance method combines structural features of the pan-caspase toxicity and thus suffers the same limitation. For the prediction of caspase toxicity, their model performed slightly worse than the classic model naive bayesian while our results showed graph-based approaches, particularly graph transformers, and consensus models generally achieved the best results. In the following, we offer a comprehensive analysis on the cross-cell-line caspase activations and cell-line-specific structural motifs that are exclusively active for a chosen cell type.

##### Chemical Structures of the Top Ten Most Broadly Active Compounds

We first analyze the structures resulting in the Caspase-3/7 activation for the largest number of cell lines. The top ten compounds, plotted in **Figure 10**, impacted a significant number, ten to six out of the fourteen cell lines tested and their structural similarity were analyzed using pairwise Tanimoto similarity. The similarity matrix in Figure 10 reveals consistently low pairwise similarity across the set, indicating highly diverse structures can result in similarly broad caspase toxicity across multiple cell lines. We further performed a clustering analysis and extracted the common structure in each cluster. The Maximum Common Substructure (MCS)-based scaffold extraction was adopted, which is a cheminformatics approach used to identify the largest shared structural core among a group of compounds. Only two pairs of compounds exhibit notable scaffold overlap: compounds 2724411 and 11057 share a 25-atom scaffold, and compounds 62770 and 443939 share a 38-atom scaffold, as highlighted in Figure 10. The remaining six compounds form singleton clusters with no significant shared substructures.

**Figure 10:**
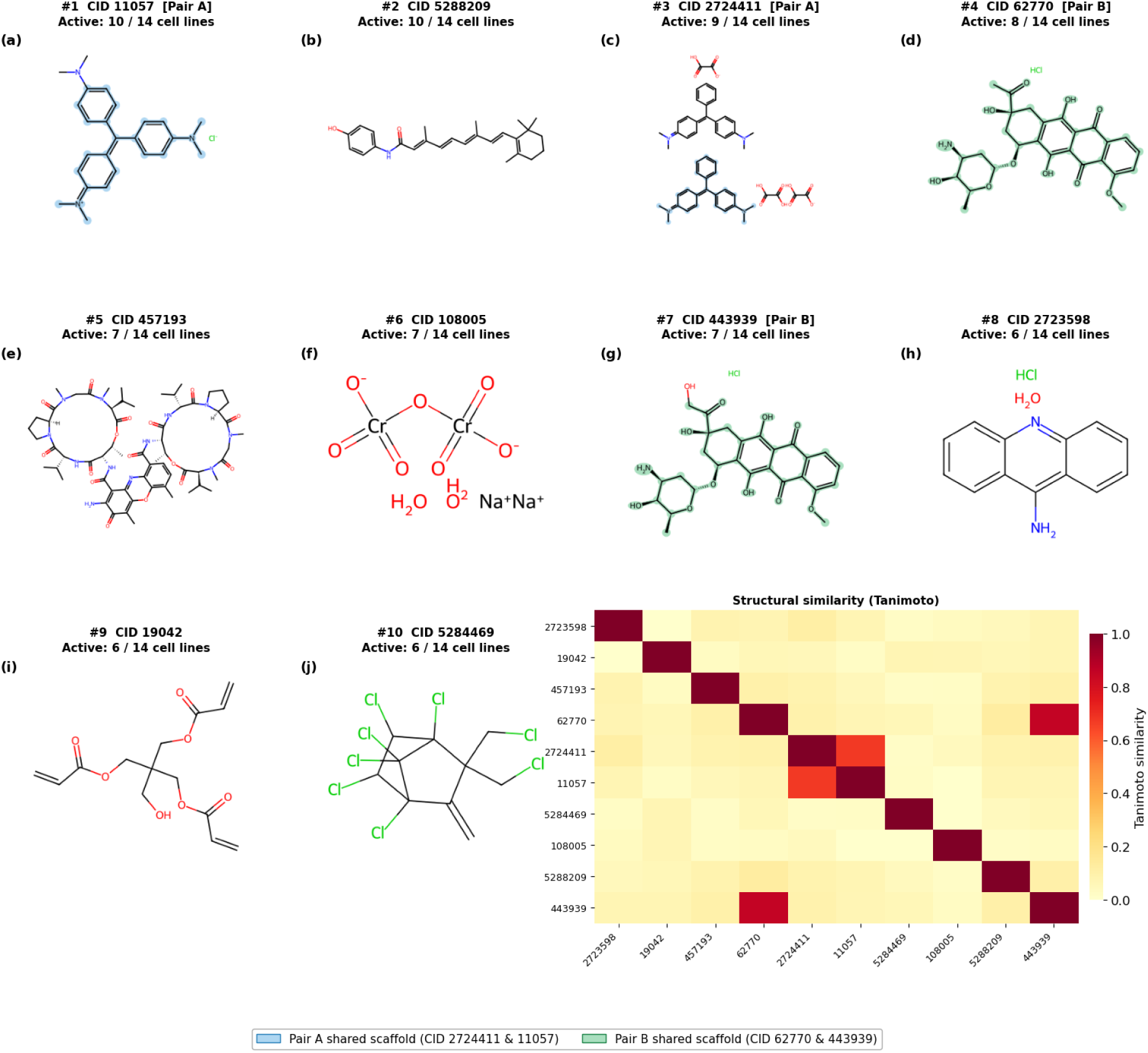
Structures of the top ten most broadly active Caspase-3/7-activating compounds, sorted by decreasing number of active cell lines. The maximum common substructures (MCS) for two pairs of compounds are identified and highlighted: Pair A (CID 2724411 and 11057, blue) and Pair B (CID 62770 and 443939, green). The remaining six compounds share no significant common substructures. Pairwise Tanimoto similarity matrix confirms low overall structural similarity across the set.

Despite this structural diversity, all compounds demonstrate activity across multiple cell lines, suggesting that Caspase-3/7 activation is not driven by a single conserved scaffold but instead may arise from diverse chemotypes associated with complex cytotoxic mechanisms.

##### Cell-Line-Specific Structural Motifs

To assess whether structural features predict cell-line selectivity, BRICS fragment enrichment was applied to each cell line. A one-sided Fisher’s exact test compared fragments enriched in compounds active in a given cell line against compounds active in other cell lines. The FDR threshold was set to *α* = 0.10 because per-cell-line foreground groups were small (*n* = 10–19 active compounds); at *α* = 0.05 the test is severely underpowered for groups of this size. Six cell lines with fewer than ten active compounds were excluded from further analysis. For the remaining eight cell lines, HepG2 (*n* = 241), CHO-K1 (*n* = 115), Jurkat (*n* = 48), and SH-SY5Y (*n* = 10) returned zero significant fragments. However, SK-N-SH (*n* = 19), H-4-II-E (*n* = 19), and HEK293 (*n* = 17) produced enriched fragments; these are illustrated in **Figure 11**, with full enrichment statistics provided in Supplementary Table S3. HEK293 was the only cell line with a distinct structural pattern: 1,1-dichloroethane (OR = *∞*, *p*_adj_ = 0.010) and chlorobenzene (OR = 10.7, *p*_adj_ = 0.022) were enriched exclusively in HEK293 and were absent from all other cell lines. SK-N-SH uniquely enriched an epoxide fragment (OR = 35.6, *p*_adj_ = 0.050). H-4-II-E uniquely enriched a tetramethylcyclohexene motif (OR = *∞*, *p*_adj_ = 0.001) and an acetaldehyde fragment (OR = 35.6, *p*_adj_ = 0.032).

**Figure 11:**
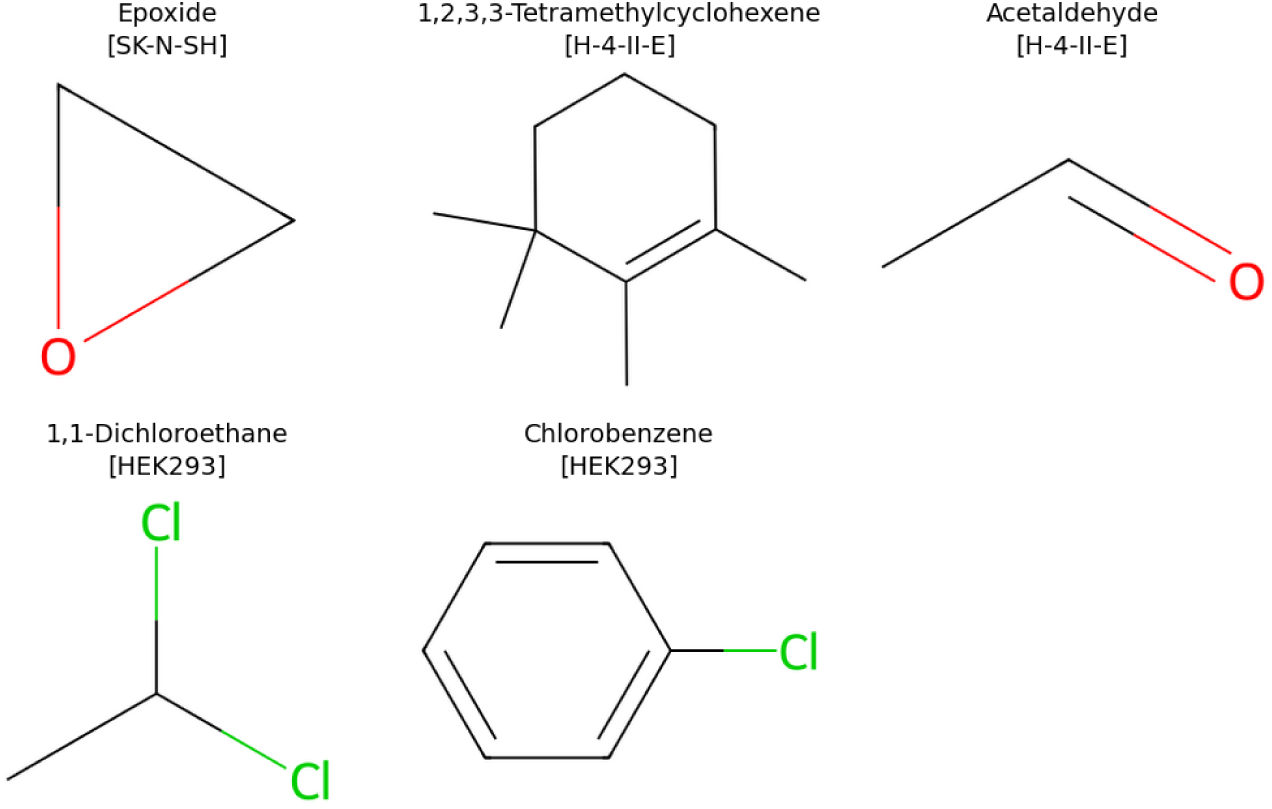
BRICS fragments significantly enriched in **cell-line-specific** active compound sets. One fragment is unique to SK-N-SH; two are unique to H-4-II-E; two are unique to HEK293. No significant fragments were found for HepG2, CHO-K1, Jurkat, or SH-SY5Y. Full enrichment statistics are in Supplementary Table S3.

The observed differences in Caspase-3/7 activation across these cell lines can be attributed to the interplay between their chemical structures and the cell-specific capacity for biotransformation. In SK-N-SH neuroblastoma cells, the presence of an epoxide functional group introduces a highly reactive electrophilic center; this feature is capable of directly forming adducts with proteins and DNA, thereby triggering apoptotic signaling. Notably, neuroblastoma cells such as SK-N-SH lack certain detoxification enzymes (including microsomal glutathione S-transferase 1, or MGST1); this deficiency impairs the cells’ ability to neutralize electrophilic intermediates, allowing these intermediates to persist and subsequently promote Caspase activation [41]. Unlike in hepatic systems, CYP enzyme-mediated bioactivation is also relatively limited in SK-N-SH cells; this implies that in this specific cell line, direct chemical reactivity specifically the epoxide structure—rather than metabolic transformation, is the primary driver of apoptosis. In contrast, the situation is markedly different in H-4-II-E cells, which are derived from hepatocytes; this distinction is primarily attributable to their robust capacity for xenobiotic metabolism. These cells express inducible CYP1A1/CYP1A2 and other Phase I metabolic enzymes, enabling them to convert lipophilic compounds—such as 1,2,3,3-tetramethylcyclohexene into reactive epoxide intermediates. Concurrently, aldehyde-containing structures (e.g., acetaldehyde) can undergo further transformation via metabolic pathways involving alcohol dehydrogenase (ADH) and CYP2E1, thereby generating electrophilic metabolites [42, 43]. It is well established that within hepatocyte systems, these reactive species are capable of activating apoptotic signaling cascades and promoting the activation of Caspase-3/7 [44]. Consequently, the robust metabolic capacity of H-4-II-E cells amplifies chemical toxicity by converting relatively inert lipophilic precursors into active electrophilic reagents; this also explains why these cells exhibit heightened sensitivity compared to non-hepatic cell lines.

Regarding HEK293 cells, it is well established that halogenated hydrocarbons disrupt cell membrane integrity and induce stress responses; these stress responses can subsequently trigger apoptosis via stress kinase pathways (e.g., JNK) and the downstream activation of Caspases [44]. In contrast, SK-N-SH cells rely more heavily on direct electrophilic damage (e.g., from epoxides) to achieve robust Caspase activation; consequently, this cell line may exhibit a relatively attenuated response to the weaker stress signals generated by halogenated compounds. Furthermore, differences across cell types—specifically regarding their apoptotic priming status (e.g., the balance of Bcl-2 family proteins) and basal redox state—also influence the threshold required to trigger Caspase activation.

#### Mitochondria Pathway and Cytotoxicity

Since the cytotoxicity in terms of the cell viability in the same assay well of MMP assay is also available, we conducted pairwise comparison of MMP activity and cell viability in **Table 9**. As shown in Table 9, 6% of compounds were active in both MMP activity and cell viability. Notably, no compounds were found that were mitochondria-inactive and viability-active, indicating that the cell viability loss does not occur independently of MMP disruption under these assay conditions.

**Table 9:**
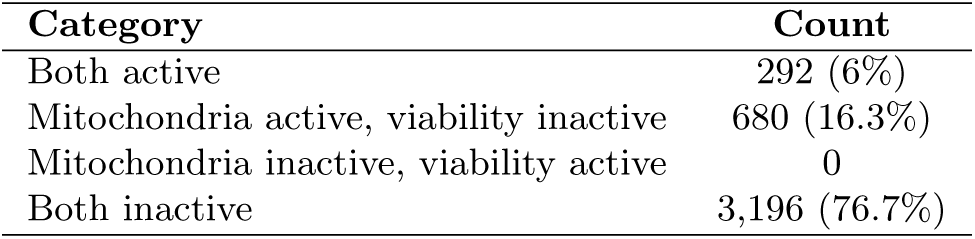
Pairwise comparison of mitochondrial activity and cell viability in HepG2 cells.

| Category | Count |
| --- | --- |
| Both active | 292 (6%) |
| Mitochondria active, viability inactive | 680 (16.3%) |
| Mitochondria inactive, viability active | 0 |
| Both inactive | 3,196 (76.7%) |

##### Structural Motifs Associated with Dual Mitochondrial and Cell Viability Activity

To identify structural features associated with simultaneous mitochondrial membrane disruption and reduced cell viability, BRICS fragment enrichment was applied to compounds active in both assays (*n* = 292) against compounds inactive in both (*n* = 3,196; one-sided Fisher’s exact test, Benjamini–Hochberg FDR *α* = 0.01). Seventeen fragments reached significance across eight structural families; these are illustrated in **Figure 12**, with full enrichment statistics provided in Supplementary Table S4.

**Figure 12:**
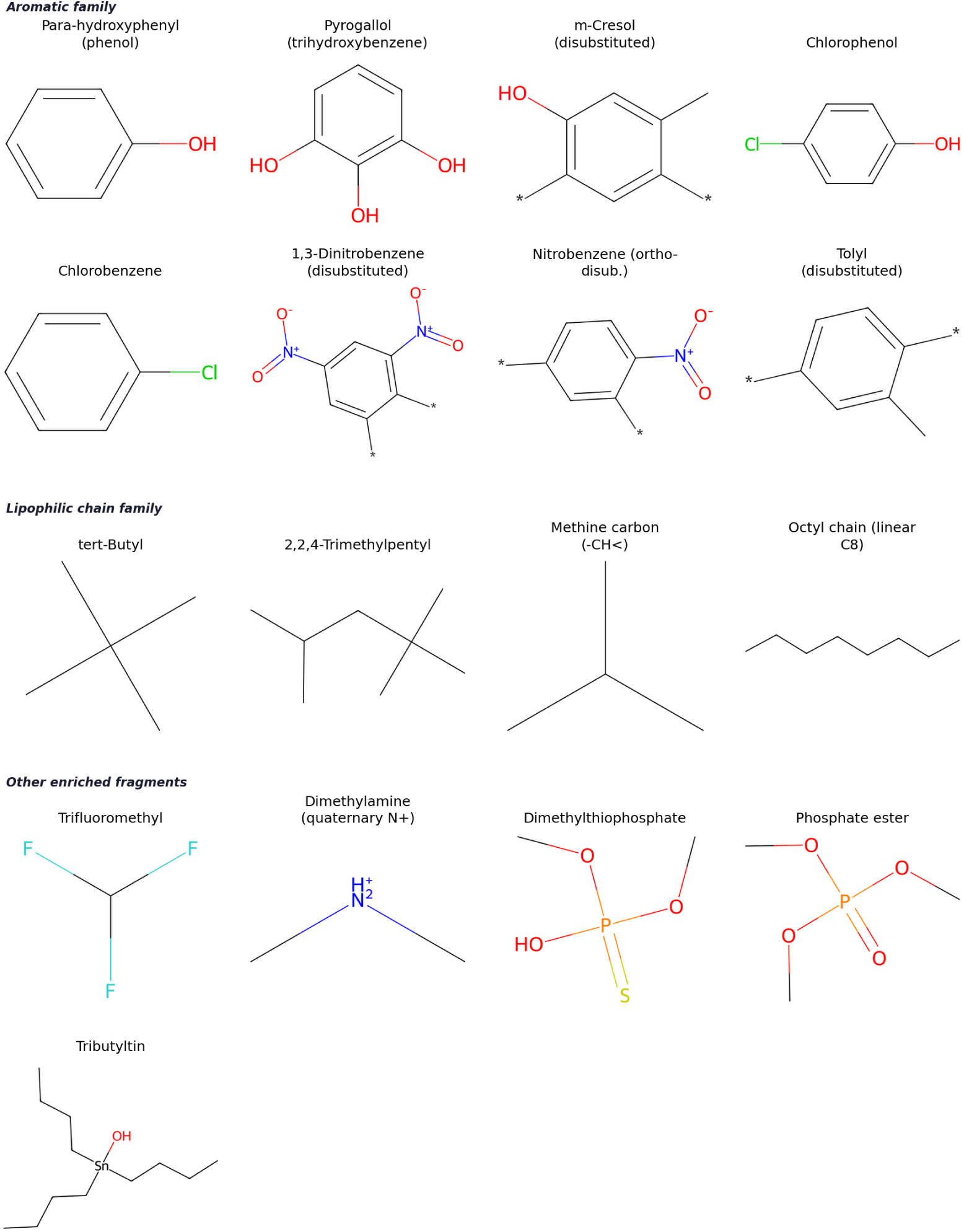
BRICS fragments significantly enriched in compounds active in both the mitochondrial membrane potential and cell viability assays versus compounds inactive in both (HepG2; *n*_active_ = 292, *n*_inactive_ = 3,196; FDR *α* = 0.01). Fragments are grouped by structural family; all 17 significant motifs are shown. Full enrichment statistics (OR, 95 % CI, *p*_adj_, prevalence) are provided in Supplementary Table S4.

The phenolic family produced four enriched motifs. Para-hydroxyphenyl (phenol) was the top-ranked fragment overall based on *p*_adj_ (*p*_adj_ = 6.0 *×* 10*^−^*^6^), followed by 4-chlorophenol (*p*_adj_ = 4.7 *×* 10*^−^*^4^), pyrogallol (*p*_adj_ = 2.9 *×* 10*^−^*^3^), and *m*-cresol (*p*_adj_ = 2.9 *×* 10*^−^*^3^), all of which are absent from the dual-inactive set. Dimethylamine was the only nitrogen-containing fragment and carried the highest point estimate among rank-2 families (*p*_adj_ = 1.7 *×* 10*^−^*^5^). Trifluoromethyl was the sole halogenated motif and ranked third overall (*p*_adj_ = 2.7 *×* 10*^−^*^5^). Among organophosphates, O,O-dimethyl thiophosphate and phosphate triester produced the largest point estimates in the analysis, consistent with known organophosphate cytotoxicity mechanisms. Tributyltin alkoxide, the sole organometallic fragment, was absent from the dual-inactive set, reflecting the well-established mitochondrial toxicity of organotin compounds. The lipophilic chain family contributed four fragments—*tert*-butyl, 2,2,4-trimethylpentyl, methine carbon, and octyl chain—indicating that branched and linear lipophilic extensions are a recurrent structural feature of dual-toxic compounds, likely facilitating membrane partitioning. Nitroaromatic and chlorinated aromatic fragments each yielded two enriched motifs at lower significance (both families *p*_adj_ *≤* 7.6 *×* 10*^−^*^3^).

##### Structural Motifs Associated with Mitochondria-Only Active Compounds

To identify structural features of compounds that disrupt mitochondrial membrane potential without reducing cell viability (*n* = 680), BRICS fragment enrichment was applied against the dual-inactive set (*n* = 3,196; one-sided Fisher’s exact test, Benjamini–Hochberg FDR *α* = 0.05, min_count = 3). Forty-eight fragments reached significance; the twelve most significant are shown in **Figure 13** with full enrichment statistics provided in Supplementary Table S5.

**Figure 13:**
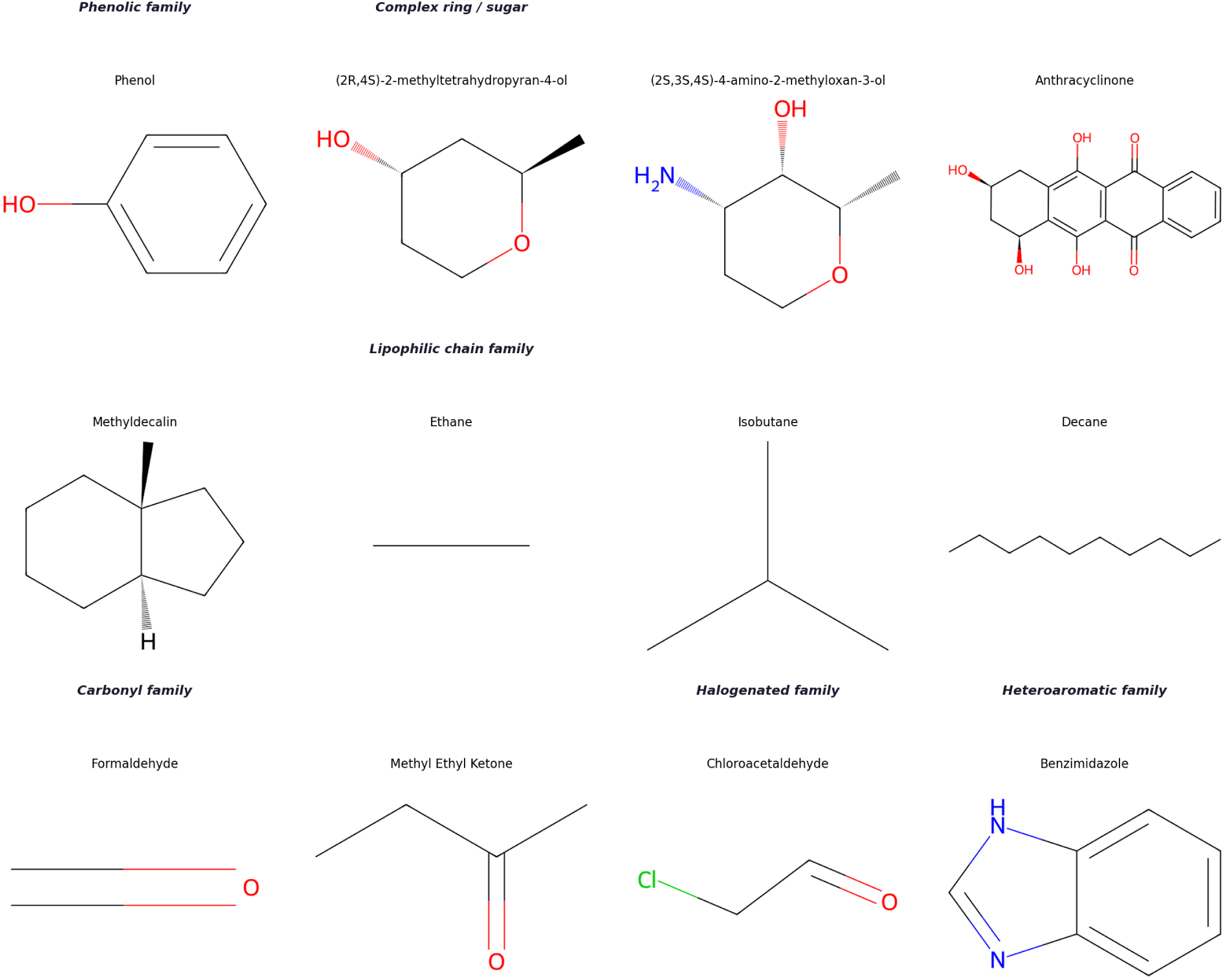
Top twelve BRICS fragments enriched in mitochondria-active, viability-inactive compounds versus dual-inactive compounds (HepG2; *n*_mito-only_ = 680, *n*_inactive_ = 3,196; FDR *α* = 0.05). Fragments are grouped by structural family; Full enrichment statistics (OR, 95 % CI, *p*_adj_, prevalence) are provided in Supplementary Table S5.

Para-hydroxyphenyl (phenol) was the top-ranked fragment and sole representative of the *phenolic family* (*p*_adj_ = 2.4 *×* 10*^−^*^5^), consistent with the known proton-uncoupling activity of phenolic compounds at mitochondrial membranes. Among *complex ring systems*, the deoxysugar (2-methyltetrahydropyran-4-ol; OR = *∞*, *p*_adj_ = 1.5 *×* 10*^−^*^4^), aminosugar, and anthracycline scaffold were entirely absent from the inactive set (OR = *∞*, *p*_adj_ = 1.7 *×* 10*^−^*^3^), reflecting the well-established mitochondrial toxicity of anthracycline-class compounds and the role of rigid polycyclic scaffolds in membrane intercalation. Methyldecalin was also enriched (OR = 28.4, *p*_adj_ = 1.7 *×* 10*^−^*^3^), consistent with the membrane-partitioning tendency of fused alicyclic rings. The *lipophilic chain family* contributed ethane (*p*_adj_ = 1.4 *×* 10*^−^*^4^), isobutane (*p*_adj_ = 1.5 *×* 10*^−^*^4^), and decane (OR = *∞*, *p*_adj_ = 1.7 *×* 10*^−^*^3^), indicating that both short-branched and long linear aliphatic chains promote MMP disruption, likely by facilitating membrane partitioning. The *carbonyl family* yielded formaldehyde (*p*_adj_ = 2.1 *×* 10*^−^*^4^) and methyl ethyl ketone (OR = *∞*, *p*_adj_ = 1.7 *×* 10*^−^*^3^), while chloroacetaldehyde was the sole *halogenated* fragment (OR = 28.4, *p*_adj_ = 1.7 *×* 10*^−^*^3^). Benzimidazole represented the *heteroaromatic family* (OR = 28.4, *p*_adj_ = 1.7 *×* 10*^−^*^3^).

## Conclusion

Caspase-3/7 is a key biomarker and the final common pathway for cells to irreversibly enter apoptosis. Simultaneous monitoring of Caspase-3/7 activity and mitochondrial membrane potential (MMP) disruption can provide a mechanistically clear series of biomarkers for evaluating compound-induced apoptotic toxicity. We propose a graph-based QSAR modeling pipeline that can take public *in vitro* and *in vivo* biological datasets as input, preprocess for machine learning training, construct feature representations, and leverage advanced graph deep learning approaches, such as graph neural networks (GNNs) and graph transformers (GTs). Graph-based approaches demonstrated outstanding performance for predicting Caspase-3/7 activation, MMP disruption, and *in vivo* FDA-approved DILI. For example in predicting human hepatotoxicity, graphormer and the full consensus model achieved an F1 score of 0.79 and an ROC-AUC of 0.69 respectively, significantly improving the previous best model on the same DILI test data. Moreover, we analyzed the mechanistic interplay between Caspase-3/7 and MMP disruption, and identified structural motifs that may be responsible for dual activation, such as fragments from the phenolic family, lipophilic chain family and complex ring systems. We also studied the impacts of cell types on Caspase-3/7 activation and extracted cell-type-specific structural motifs that are exclusively active for a chosen cell type.

For future research directions, we will enhance our graph-based QSAR modeling pipeline by feeding large-scale literature-derived toxicological data. Large language model (LLM)- and agentic AI-driven data mining holds great potential to automatically extract large-scale toxicological knowledge from rapidly expanding scientific literature. Moreover, exploring the use of advanced LLMs for compound-induced toxicity prediction and conducting systematic comparisons with our graph-based QSAR approaches represent promising avenues for future work.

## Supporting information

The following files are available free of charge.

- Supplementary Methods (PDF)
- Supplementary Tables and Figures (PDF)

## Data and Code Availability

We have released the Python code for our graph-based QSAR modeling pipeline and all the datasets used in this study, which are publicly available on GitHub at https://github.com/yaminichitikela/qsar_project_dili.

## References

(1) Bursch, W.; Karwan, A.; Mayer, M.; Dornetshuber, J.; Fröhwein, U.; Schulte-Hermann, R.; Fazi, B.; Di Sano, F.; Piredda, L.; Piacentini, M., et al. Cell death and autophagy: cytokines, drugs, and nutritional factors. Toxicology 2008, 254, 147–157.

(2) Gogvadze, V.; Orrenius, S. Mitochondrial regulation of apoptotic cell death. Chemico-biological interactions 2006, 163, 4–14.

(3) Inglese, J.; Auld, D. S.; Jadhav, A.; Johnson, R. L.; Simeonov, A.; Yasgar, A.; Zheng, W.; Austin, C. P. Quantitative high-throughput screening: a titration-based approach that efficiently identifies biological activities in large chemical libraries. Proceedings of the National Academy of Sciences 2006, 103, 11473–11478.

(4) Xia, M.; Huang, R.; Witt, K. L.; Southall, N.; Fostel, J.; Cho, M.-H.; Jadhav, A.; Smith, C. S.; Inglese, J.; Portier, C. J.; Tice, R. R.; Austin, C. P. Compound cytotoxicity profiling using quantitative high-throughput screening. Environmental Health Perspectives 2008, 116, 284–291.

(5) Attene-Ramos, M. S.; Miller, N.; Huang, R.; Michael, S.; Itkin, M.; Kavlock, R. J.; Austin, C. P.; Shinn, P.; Simeonov, A.; Tice, R. R.; Xia, M. The Tox21 robotic platform for the assessment of environmental chemicals—from vision to reality. Drug Discovery Today 2013, 18, 716–723.

(6) Wang, Y.; Suzek, T.; Zhang, J.; Wang, J.; He, S.; Cheng, T.; Shoemaker, B. A.; Gindulyte, A.; Bryant, S. H. PubChem BioAssay: 2014 update. Nucleic Acids Research 2014, 42, D1075–D1082.

(7) Hansch, C.; Fujita, T. *ρ*–*σ*–*π* analysis: a method for the correlation of biological activity and chemical structure. Journal of the American Chemical Society 1964, 86, 1616–1626.

(8) Tropsha, A. Best practices for QSAR model development, validation, and exploitation. Molecular Informatics 2010, 29, 476–488.

(9) Huang, R.; Xia, M.; Sakamuru, S.; Zhao, J.; Shahane, S. A.; Attene-Ramos, M.; Zhao, T.; Austin, C. P.; Simeonov, A. Modelling the Tox21 10K chemical profiles for in vivo toxicity prediction and mechanism characterization. Nature Communications 2016, 7, 10425.

(10) Mayr, A.; Klambauer, G.; Unterthiner, T.; Hochreiter, S. DeepTox: toxicity prediction using deep learning. Frontiers in Environmental Science 2016, 3, 80.

(11) Kim, M. T.; Huang, R.; Sedykh, A.; Wang, W.; Xia, M.; Zhu, H. Mechanism profiling of hepatotoxicity caused by oxidative stress using antioxidant response element reporter gene assay models and big data. Environmental health perspectives 2015, 124, 634.

(12) Huang, R.; Southall, N.; Xia, M.; Cho, M.-H.; Jadhav, A.; Nguyen, D.-T.; Inglese, J.; Tice, R. R.; Austin, C. P. Weighted feature significance: a simple, interpretable model of compound toxicity based on the statistical enrichment of structural features. Toxicological Sciences 2009, 112, 385–393.

(13) Gilmer, J.; Schoenholz, S. S.; Riley, P. F.; Vinyals, O.; Dahl, G. E. In *Proceedings of the 34th International Conference on Machine Learning*, PMLR: 2017; Vol. 70, pp 1263–1272.

(14) Rogers, D.; Hahn, M. Extended-connectivity fingerprints. Journal of chemical information and modeling 2010, 50, 742–754.

(15) Vaswani, A.; Shazeer, N.; Parmar, N.; Uszkoreit, J.; Jones, L.; Gomez, A. N.; Kaiser, Ł.; Polosukhin, I. Attention is all you need. Advances in neural information processing systems 2017, 30.

(16) Masters, D.; Dean, J.; Klaser, K.; Li, Z.; Maddrell-Mander, S.; Sanders, A.; Helal, H.; Beker, D.; Rampášek, L.; Beaini, D. Gps++: An optimised hybrid mpnn/transformer for molecular property prediction. arXiv preprint arXiv:2212.02229 2022.

(17) Müller, L.; Galkin, M.; Morris, C.; Rampášek, L. Attending to graph transformers. arXiv preprint arXiv:2302.04181 2023.

(18) Thai Kim, M.; Huang, R.; Sedykh, A.; Wang, W.; Xia, M.; Zhu, H. Mechanism Profiling of Hepatotoxicity Caused by Oxidative Stress Using Antioxidant Response Element Reporter Gene Assay Models and Big Data. Environmental Health Perspectives 2016, 124, DOI: 10.1289/ehp.1509763.

(19) Seal, S.; Williams, D.; Hosseini-Gerami, L.; Mahale, M.; Carpenter, A. E.; Spjuth, O.; Bender, A. Improved detection of drug-induced liver injury by integrating predicted in vivo and in vitro data. Chemical research in toxicology 2024, 37, 1290–1305.

(20) ATCC CHO-K1 cell line (CCL-61), American Type Culture Collection, Available at: https://www.atcc.org/products/ccl-61, 2024.

(21) Landrum, G. et al. RDKit: Open-source cheminformatics, 2006.

(22) Breiman, L. Random Forests. Machine Learning 2001, 45, 5–32.

(23) Cortes, C.; Vapnik, V. Support-Vector Networks. Machine Learning 1995, 20, 273–297.

(24) Chen, T.; Guestrin, C. In Proceedings of the 22nd ACM SIGKDD International Conference on Knowledge Discovery and Data Mining, ACM: 2016, pp 785–794.

(25) Kipf, T. N.; Welling, M. Semi-supervised classification with graph convolutional networks. arXiv preprint arXiv:1609.02907 2016.

(26) Veličković, P.; Cucurull, G.; Casanova, A.; Romero, A.; Lio, P.; Bengio, Y. Graph attention networks. arXiv preprint arXiv:1710.10903 2017.

(27) Xu, K.; Hu, W.; Leskovec, J.; Jegelka, S. How powerful are graph neural networks? arXiv preprint arXiv:1810.00826 2018.

(28) Hamilton, W.; Ying, Z.; Leskovec, J. Inductive representation learning on large graphs. Advances in neural information processing systems 2017, 30.

(29) Fey, M.; Lenssen, J. E. Fast graph representation learning with PyTorch Geometric. arXiv preprint arXiv:1903.02428 2019.

(30) Ying, C.; Cai, T.; Luo, S.; Zheng, S.; Ke, G.; He, D.; Shen, Y.; Liu, T.-Y. Do transformers really perform badly for graph representation? Advances in neural information processing systems 2021, 34, 28877–28888.

(31) Rampášek, L.; Galkin, M.; Dwivedi, V. P.; Luu, A. T.; Wolf, G.; Beaini, D. Recipe for a general, powerful, scalable graph transformer. Advances in Neural Information Processing Systems 2022, 35, 14501–14515.

(32) Thakkar, S.; Li, T.; Liu, Z.; Wu, L.; Roberts, R.; Tong, W. Drug-induced liver injury severity and toxicity (DILIst): binary classification of 1279 drugs by human hepatotoxicity. Drug discovery today 2020, 25, 201–208.

(33) Chen, M.; Suzuki, A.; Thakkar, S.; Yu, K.; Hu, C.; Tong, W. DILIrank: the largest reference drug list ranked by the risk for developing drug-induced liver injury in humans. Drug discovery today 2016, 21, 648–653.

(34) Yuan, H.; Yu, H.; Wang, J.; Li, K.; Ji, S. In Proceedings of the 38th International Conference on Machine Learning, PMLR: 2021; Vol. 139, pp 12241–12252.

(35) Seal, S.; Williams, D. P.; Hosseini-Gerami, L.; Mahale, M.; Carpenter, A. E.; Spjuth, O.; Bender, A. Improved Detection of Drug-Induced Liver Injury by Integrating Predicted In Vivo and In Vitro Data. Chemical Research in Toxicology 2024, 37, 1290–1305.

(36) Degen, J.; Wegscheid-Gerlach, C.; Zaliani, A.; Rarey, M. On the art of compiling and using’drug-like’chemical fragment spaces. ChemMedChem 2008, 3, 1503.

(37) Džajić, I.; Tomasic, T.; Pardo, L. A.; Peterlin Masic, L.; Cotman, A. E. Lipophilic Cations as Mitochondria-Targeting Moieties: Recent Progress and Design Principles for Medicinal Chemistry. Journal of medicinal chemistry 2025, 68, 23690–23704.

(38) Trnka, J.; Elkalaf, M.; Anděl, M. Lipophilic triphenylphosphonium cations inhibit mitochondrial electron transport chain and induce mitochondrial proton leak. PloS one 2015, 10, e0121837.

(39) Fernandes, C.; Videira, A. J.; Veloso, C. D.; Benfeito, S.; Soares, P.; Martins, J. D.; Gonçalves, B.; Duarte, J. F.; Santos, A. M.; Oliveira, P. J., et al. Cytotoxicity and mitochondrial effects of phenolic and quinone-based mitochondria-targeted and untargeted antioxidants on human neuronal and hepatic cell lines: a comparative analysis. Biomolecules 2021, 11, 1605.

(40) Finichiu, P. G.; James, A. M.; Larsen, L.; Smith, R. A.; Murphy, M. P. Mitochondrial accumulation of a lipophilic cation conjugated to an ionisable group depends on membrane potential, pH gradient and p K a: implications for the design of mitochondrial probes and therapies. Journal of bioenergetics and biomembranes 2013, 45, 165–173.

(41) Kelner, M. J.; Diccianni, M. B.; Alice, L. Y.; Rutherford, M. R.; Estes, L. A.; Morgenstern, R. Absence of MGST1 mRNA and protein expression in human neuroblastoma cell lines and primary tissue. Free Radical Biology and Medicine 2014, 69, 167–171.

(42) Kaisarevic, S.; Dakic, V.; Hrubik, J.; Glisic, B.; Lübcke-von Varel, U.; Pogrmic-Majkic, K.; Fa, S.; Teodorovic, I.; Brack, W.; Kovacevic, R. Differential expression of CYP1A1 and CYP1A2 genes in H4IIE rat hepatoma cells exposed to TCDD and PAHs. Environmental toxicology and pharmacology 2015, 39, 358–368.

(43) Marković Filipović, J.; Miler, M.; Kojić, D.; Karan, J.; Ivelja, I.; Čukuranović Kokoris, J.; Matavulj, M. Effect of acrylamide treatment on Cyp2e1 expression and redox status in rat hepatocytes. International Journal of Molecular Sciences 2022, 23, 6062.

(44) Michels, G.; Wätjen, W.; Weber, N.; Niering, P.; Chovolou, Y.; Kampkötter, A.; Proksch, P.; Kahl, R. Resveratrol induces apoptotic cell death in rat H4IIE hepatoma cells but necrosis in C6 glioma cells. Toxicology 2006, 225, 173–182.

