## Supplemental File 1 for "A Graph-based QSAR Modeling Pipeline for Predicting In vitro PubChem Assays and In vivo Human hepatotoxicity: Mechanistic Analysis of Caspase-3/7 Activation"

This PDF file includes:

- Table S1: QSAR Model Performance in the HepG2 Caspase-3/7 dataset
- Table S2: QSAR Model Performance in the CHO-K1 Caspase-3/7 dataset
- Table S3: Cell-line-specific Structural Motifs
- Table S4: Structural Motifs Associated with Dual Mitochondrial and Cell Viability Activity.
- Table S5: Structural Motifs Associated with Mitochondria-active, Viability-inactive Compounds vs Dual-inactive Compounds
- Figure S1: ROC-AUC and PR-AUC curves in the HepG2 Caspase-3/7 dataset
- Figure S2: ROC-AUC and PR-AUC curves in the CHO-K1 Caspase-3/7 dataset

Table S1: Test set performance of all models at decision thresholds CPT-1 (0.50) and CPT-2 (0.80) for the HepG2 Caspase-3/7 dataset. Sens = sensitivity (%); Spec = specificity (%); CCR = correct classification rate (%). Bal-Acc = balanced accuracy (%).

| Model | CPT | Sens | Spec | CCR | ROC-AUC | PR-AUC | MCC | Bal-Acc | F1 |
|---|---|---|---|---|---|---|---|---|---|
| *Classical Machine Learning* | | | | | | | | | |
| Random Forest | 0.50 | 56.0 | 78.6 | 67.3 | **0.7956** | **0.1682** | 0.1493 | 67.3 | 0.1455 |
| | 0.80 | 8.0 | 99.3 | 53.7 | **0.7956** | **0.1682** | 0.1363 | 53.7 | 0.1250 |
| SVM | 0.50 | 72.0 | 68.1 | 70.0 | 0.7769 | 0.1679 | 0.1534 | 70.0 | 0.1324 |
| | 0.80 | 6.0 | **99.9** | 52.9 | 0.7769 | 0.1679 | 0.1825 | 52.9 | 0.1091 |
| XGBoost | 0.50 | 60.0 | 63.5 | 61.7 | 0.6946 | 0.0941 | 0.0875 | 61.7 | 0.0993 |
| | 0.80 | 32.0 | 89.4 | 60.7 | 0.6946 | 0.0941 | 0.1219 | 60.7 | 0.1468 |
| *Graph Neural Networks* | | | | | | | | | |
| GCN | 0.50 | 68.0 | 72.2 | 70.1 | 0.7622 | 0.0925 | 0.1599 | 70.1 | 0.1411 |
| | 0.80 | 28.0 | 93.0 | 60.5 | 0.7622 | 0.0925 | 0.1414 | 60.5 | 0.1697 |
| GAT | 0.50 | 74.0 | 61.9 | 67.9 | 0.7449 | 0.0886 | 0.1324 | 67.9 | 0.1167 |
| | 0.80 | 46.0 | 85.8 | 65.9 | 0.7449 | 0.0886 | 0.1593 | 65.9 | 0.1661 |
| GIN | 0.50 | 74.0 | 69.7 | 71.9 | 0.7465 | 0.0940 | 0.1695 | 71.9 | 0.1420 |
| | 0.80 | 48.0 | 85.4 | 66.7 | 0.7465 | 0.0940 | 0.1651 | 66.7 | 0.1690 |
| GraphSAGE | 0.50 | 68.0 | 64.6 | 66.3 | 0.7275 | 0.0851 | 0.1221 | 66.3 | 0.1149 |
| | 0.80 | 32.0 | 90.9 | 61.5 | 0.7275 | 0.0851 | 0.1389 | 61.5 | 0.1633 |
| *Graph Transformers* | | | | | | | | | |
| Graphormer | 0.50 | 76.0 | 36.1 | 56.1 | 0.6164 | 0.0485 | 0.0457 | 56.1 | 0.0757 |
| | 0.80 | 14.0 | 91.6 | 52.8 | 0.6164 | 0.0485 | 0.0363 | 52.8 | 0.0791 |
| GraphGPS | 0.50 | 64.0 | 67.7 | 65.9 | 0.7412 | 0.0956 | 0.1214 | 65.9 | 0.1174 |
| | 0.80 | 32.0 | 92.3 | 62.1 | 0.7412 | 0.0956 | 0.1565 | 62.1 | 0.1808 |
| *Consensus Ensembles* | | | | | | | | | |
| Consensus-Classical | 0.50 | 68.0 | 69.9 | 69.0 | 0.7684 | 0.1374 | 0.1476 | 69.0 | 0.1320 |
| | 0.80 | 14.0 | 98.4 | 56.2 | 0.7684 | 0.1374 | 0.1589 | 56.2 | 0.1750 |
| Consensus-GNN | 0.50 | 70.0 | 68.8 | 69.4 | 0.7606 | 0.0995 | 0.1493 | 69.4 | 0.1313 |
| | 0.80 | 36.0 | 90.9 | 63.5 | 0.7606 | 0.0995 | 0.1622 | 63.5 | **0.1818** |
| Consensus-GT | 0.50 | 68.0 | 60.7 | 64.3 | 0.7123 | 0.0869 | 0.1055 | 64.3 | 0.1049 |
| | 0.80 | 14.0 | 96.8 | 55.4 | 0.7123 | 0.0869 | 0.1049 | 55.4 | 0.1359 |
| Consensus | 0.50 | **78.0** | 69.3 | **73.7** | 0.7910 | 0.1371 | **0.1826** | **73.7** | 0.1474 |
| | 0.80 | 10.0 | 98.2 | 54.1 | 0.7910 | 0.1371 | 0.1033 | 54.1 | 0.1235 |

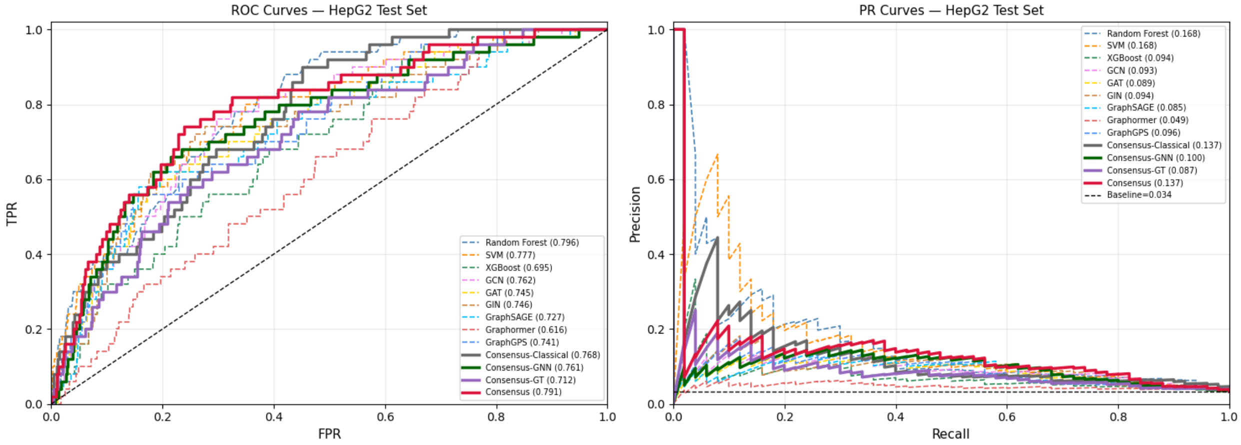


Figure S1: ROC-AUC and PR-AUC curves for all thirteen models on the HepG2 Caspase-3/7 dataset

Table S2: Test set performance of all models at decision thresholds CPT-1 (0.50) and CPT-2 (0.80) for the CHO-K1 Caspase-3/7 dataset. Sens = sensitivity (%); Spec = specificity (%); CCR = correct classification rate (%). Bal-Acc = balanced accuracy (%).

| **Model** | **CPT** | **Sens** | **Spec** | **CCR** | **ROC-AUC** | **PR-AUC** | **MCC** | **Bal-Acc** | **F1** |
|---|---|---|---|---|---|---|---|---|---|
| *Classical Machine Learning* | | | | | | | | | |
| Random Forest | 0.50 | 56.5 | 81.5 | 69.0 | 0.7625 | 0.1258 | 0.1193 | 69.0 | 0.0847 |
| | 0.80 | 4.3 | **99.9** | 52.1 | 0.7625 | 0.1258 | 0.1441 | 52.1 | 0.0800 |
| SVM | 0.50 | 69.6 | 63.0 | 66.3 | 0.7837 | 0.1223 | 0.0831 | 66.3 | 0.0552 |
| | 0.80 | 26.1 | 98.1 | 62.1 | 0.7837 | 0.1223 | 0.1996 | 62.1 | **0.2105** |
| XGBoost | 0.50 | 65.2 | 62.5 | 63.9 | 0.6670 | 0.0783 | 0.0705 | 63.9 | 0.0511 |
| | 0.80 | 52.2 | 83.1 | 67.7 | 0.6670 | 0.0783 | 0.1149 | 67.7 | 0.0851 |
| *Graph Neural Networks* | | | | | | | | | |
| GCN | 0.50 | 69.6 | 64.6 | 67.1 | 0.7212 | 0.0303 | 0.0879 | 67.1 | 0.0575 |
| | 0.80 | 21.7 | 86.1 | 53.9 | 0.7212 | 0.0303 | 0.0277 | 53.9 | 0.0431 |
| GAT | 0.50 | 47.8 | 70.2 | 59.0 | 0.7179 | 0.0397 | 0.0483 | 59.0 | 0.0467 |
| | 0.80 | 26.1 | 90.2 | 58.2 | 0.7179 | 0.0397 | 0.0671 | 58.2 | 0.0698 |
| GIN | 0.50 | 60.9 | 66.5 | 63.7 | 0.7427 | 0.0406 | 0.0714 | 63.7 | 0.0531 |
| | 0.80 | 39.1 | 78.6 | 58.8 | 0.7427 | 0.0406 | 0.0529 | 58.8 | 0.0520 |
| GraphSAGE | 0.50 | 60.9 | 71.7 | 66.3 | 0.7105 | 0.0391 | 0.0886 | 66.3 | 0.0619 |
| | 0.80 | 34.8 | 85.3 | 60.0 | 0.7105 | 0.0391 | 0.0695 | 60.0 | 0.0650 |
| *Graph Transformers* | | | | | | | | | |
| Graphormer | 0.50 | 69.6 | 67.8 | 68.7 | 0.7728 | 0.0760 | 0.0981 | 68.7 | 0.0626 |
| | 0.80 | 26.1 | 95.6 | 60.9 | 0.7728 | 0.0760 | 0.1265 | 60.9 | 0.1290 |
| GraphGPS | 0.50 | **82.6** | 62.2 | 72.4 | 0.7897 | 0.0623 | 0.1137 | 72.4 | 0.0639 |
| | 0.80 | 73.9 | 74.2 | 74.1 | 0.7897 | 0.0623 | 0.1347 | 74.1 | 0.0815 |
| *Consensus Ensembles* | | | | | | | | | |
| Consensus-Classical | 0.50 | 65.2 | 68.4 | 66.8 | 0.7379 | 0.1430 | 0.0888 | 66.8 | 0.0599 |
| | 0.80 | 21.7 | 98.5 | 60.1 | 0.7379 | **0.1430** | 0.1870 | 60.1 | 0.2000 |
| Consensus-GNN | 0.50 | 56.5 | 69.9 | 63.2 | 0.7336 | 0.0380 | 0.0707 | 63.2 | 0.0545 |
| | 0.80 | 30.4 | 89.5 | 60.0 | 0.7336 | 0.0380 | 0.0796 | 60.0 | 0.0765 |
| Consensus-GT | 0.50 | **82.6** | 65.8 | **74.2** | **0.8319** | 0.1230 | 0.1252 | **74.2** | 0.0700 |
| | 0.80 | 47.8 | 94.1 | 71.0 | **0.8319** | 0.1230 | **0.2096** | 71.0 | 0.1833 |
| Consensus | 0.50 | 73.9 | 70.8 | 72.3 | 0.7958 | **0.1636** | 0.1204 | 72.3 | 0.0726 |
| | 0.80 | 21.7 | 98.4 | 60.1 | 0.7958 | 0.1636 | 0.1831 | 60.1 | 0.1961 |

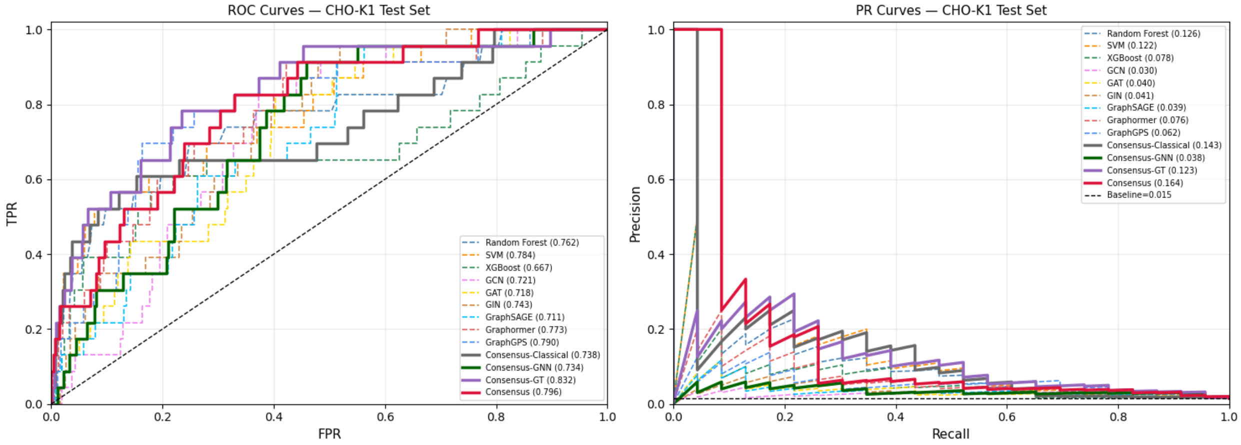


Figure S2: ROC-AUC and PR-AUC curves for all thirteen models on the CHO-K1 Caspase-3/7 dataset

Table S3: Significantly enriched BRICS fragments per cell line among the 323 Caspase-active compounds (Fisher's exact test, BH-FDR $\alpha = 0.10$, minimum foreground count = 2). OR = $\infty$ denotes fragments absent from the background set. $n_a/N_a$ = count in foreground / foreground total; $n_b/N_b$ = count in background / background total.

| **Fragment name** | **Structure** | **Cell line** | **OR** | $p_{\mathbf{adj}}$ | $n_a/N_a$ (%) | $n_b/N_b$ (%) |
|---|---|---|---|---|---|---|
| *SK-N-SH only* ($N_a = 19$, $N_b = 304$) | | | | | | |
| Epoxide | 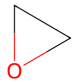 | SK-N-SH | 35.6 | 0.05 | 10.5 | 0.3 |
| *H-4-II-E only* ($N_a = 19$, $N_b = 304$) | | | | | | |
| 1,2,3,3-Tetramethylcyclohexene | 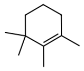 | H-4-II-E | $\infty$ | 0.001 | 15.8 | 0 |
| Acetaldehyde | 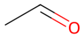 | H-4-II-E | 35.6 | 0.032 | 10.5 | 0.3 |
| *HEK293 only* ($N_a = 17$, $N_b = 306$) | | | | | | |
| 1,1-Dichloroethane | 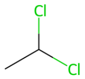 | HEK293 | $\infty$ | 0.01 | 11.8 | 0 |
| Chlorobenzene | 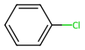 | HEK293 | 10.7 | 0.022 | 17.6 | 2.0 |

Table S4: All 17 BRICS fragments significantly enriched in dual-active (mitochondrial membrane potential + cell viability, HepG2) versus dual-inactive compounds ($n_{\text{active}} = 292$, $n_{\text{inactive}} = 3{,}196$; min. prevalence $\geq 3$ in active set, OR $\geq 2$, Benjamini–Hochberg FDR $\alpha = 0.01$). OR and 95 % CI are conditional maximum-likelihood estimates (Fisher exact); OR $= \infty$ denotes fragments absent from the dual-inactive set. % active and % inactive denote percentages of dual-active and dual-inactive compounds with a fragment respectively. *(Part 1 of 2)*

| **Fragment name** | **Structure** | **OR (95 % CI)** | $p_{\text{adj}}$ | **% active vs % inactive** |
|---|---|---|---|---|
| *Phenolic family* | | | | |
| Para-hydroxyphenyl (phenol) | 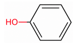 | 5.9 (3.1–10.9) | $6.0 \times 10^{-6}$ | 6.16% vs 1.10% |
| 4-Chlorophenol | 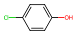 | $\infty$ (7.3, $\infty$) | $4.7 \times 10^{-4}$ | 1.37% vs 0% |
| Pyrogallol (benzene-1,2,3-triol) | 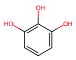 | $\infty$ (4.5, $\infty$) | $2.9 \times 10^{-3}$ | 1.03% vs 0% |
| *m*-Cresol (3-methylphenol) | 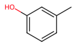 | $\infty$ (4.5, $\infty$) | $2.9 \times 10^{-3}$ | 1.03% vs 0% |
| *Lipophilic chain family* | | | | |
| *tert*-Butyl (branched C4) | 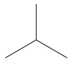 | 7.2 (3.1–15.7) | $4.6 \times 10^{-5}$ | 4.11% vs 0.59% |
| 2,2,4-Trimethylpentyl (branched C8) | 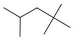 | 44.3 (4.4–2186) | $1.7 \times 10^{-3}$ | 1.37% vs 0.03% |
| Methine carbon (−CH<) | 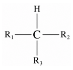 | 9.2 (2.2–36.6) | $5.1 \times 10^{-3}$ | 1.71% vs 0.19% |
| Octyl chain (linear C8) | | 7.9 (2.0–29.2) | $7.6 \times 10^{-3}$ | 1.71% vs 0.22% |
| *Nitroaromatic family* | | | | |
| 1,3-Dinitrophenyl | 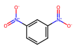 | $\infty$ (4.5, $\infty$) | $2.9 \times 10^{-3}$ | 1.03% vs 0% |
| Ortho-disubstituted nitrophenyl | 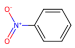 | 33.1 (2.6–1742) | $7.6 \times 10^{-3}$ | 1.03% vs 0.03% |

Table S4: *(Part 2 of 2)*

| **Fragment name** | **Structure** | **OR (95 % CI)** | $p_{\mathrm{adj}}$ | **% active vs % inactive** |
|---|---|---|---|---|
| *Chlorinated aromatic family* | | | | |
| Para-chlorophenyl | 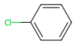 | 4.2 (1.8–9.0) | $3.0 \times 10^{-3}$ | 3.42% vs 0.85% |
| 2-Methyl-1,4-disubstituted phenyl | 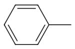 | 33.1 (2.6–1742) | $7.6 \times 10^{-3}$ | 1.03% vs 0.03% |
| *Halogenated* | | | | |
| Trifluoromethyl | 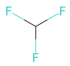 | 6.4 (3.0–12.9) | $2.7 \times 10^{-5}$ | 4.79% vs 0.78% |
| *Nitrogen-containing* | | | | |
| Dimethylamine (tertiary amine) | 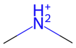 | 16.9 (5.3–58.0) | $1.7 \times 10^{-5}$ | 3.08% vs 0.19% |
| *Organophosphate family* | | | | |
| O,O-Dimethyl thiophosphate | 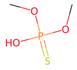 | 55.5 (6.2–2633) | $2.6 \times 10^{-4}$ | 1.71% vs 0.03% |
| Phosphate triester | 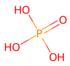 | 18.5 (3.6–120) | $1.5 \times 10^{-3}$ | 1.71% vs 0.09% |
| *Organometallic* | | | | |
| Tributyltin alkoxide† | 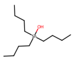 | $\infty$ (4.5, $\infty$) | $2.9 \times 10^{-3}$ | 1.03% vs 0% |

Table S5: Top twelve BRICS fragments significantly enriched in mitochondria-active, viability-inactive compounds versus dual-inactive compounds ($n_{\text{mito-only}} = 680$, $n_{\text{inactive}} = 3{,}196$; min. prevalence $\geq 3$ in foreground, Benjamini–Hochberg FDR $\alpha = 0.05$). OR and 95 % CI are conditional maximum-likelihood estimates (Fisher exact); OR $= \infty$ denotes fragments absent from the dual-inactive set. % mito-only and % inactive denote percentages of mitochondria-active, viability-inactive and dual-inactive compounds with a fragment respectively.

| **Fragment name** | **Structure** | **OR (95 % CI)** | $p_{\text{adj}}$ | **% mito-only vs % inactive** |
|---|---|---|---|---|
| *Phenolic family* | | | | |
| Para-hydroxyphenyl (phenol) | 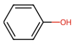 | 4.0 (2.4–6.8) | $2.4 \times 10^{-5}$ | 4.26% vs 1.10% |
| *Complex ring / sugar family* | | | | |
| 2-Methyltetrahydropyran-4-ol | 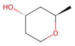 | $\infty$ (6.8, $\infty$) | $1.5 \times 10^{-4}$ | 1.03% vs 0% |
| 4-Amino-2-methyloxan-3-ol | 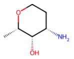 | $\infty$ (4.3, $\infty$) | $1.7 \times 10^{-3}$ | 0.74% vs 0% |
| Anthracyclinone | 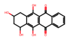 | $\infty$ (4.3, $\infty$) | $1.7 \times 10^{-3}$ | 0.74% vs 0% |
| Methyldecalin | 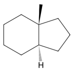 | 28.4 (3.4, $\infty$) | $1.7 \times 10^{-3}$ | 0.88% vs 0.03% |
| *Lipophilic chain family* | | | | |
| Ethane | | 7.2 (3.0–18.0) | $1.4 \times 10^{-4}$ | 2.21% vs 0.31% |
| Isobutane | 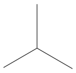 | 4.8 (2.4–9.6) | $1.5 \times 10^{-4}$ | 2.79% vs 0.59% |
| Decane | | $\infty$ (4.3, $\infty$) | $1.7 \times 10^{-3}$ | 0.74% vs 0% |
| *Carbonyl family* | | | | |
| Formaldehyde | | 5.5 (2.5–12.2) | $2.1 \times 10^{-4}$ | 2.35% vs 0.44% |
| Methyl ethyl ketone | 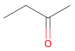 | $\infty$ (4.3, $\infty$) | $1.7 \times 10^{-3}$ | 0.74% vs 0% |
| *Halogenated family* | | | | |
| Chloroacetaldehyde | 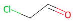 | 28.4 (3.4, $\infty$) | $1.7 \times 10^{-3}$ | 0.88% vs 0.03% |
| *Heteroaromatic family* | | | | |
| Benzimidazole | 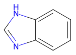 | 28.4 (3.4, $\infty$) | $1.7 \times 10^{-3}$ | 0.88% vs 0.03% |
