## Supplemental File 2 for "A Graph-based QSAR Modeling Pipeline for Predicting In vitro PubChem Assays and In vivo Human hepatotoxicity: Mechanistic Analysis of Caspase-3/7 Activation"

This PDF file includes:

- Cell Line Descriptions
- Graph Transformer Architectures
- Hyperparameter Settings
- Class Balancing: Undersampling Ratio Comparison
- SI References

### Cell Line Descriptions

Brief descriptions of each cell line are provided below, with ATCC accession numbers verified against the ATCC reference database.

- **HepG2** (ATCC HB-8065) – A human hepatocellular carcinoma cell line isolated from a 15-year-old male patient, widely used to model hepatotoxicity. It retains many liver-specific metabolic functions relevant for assessing compound-induced liver injury [1].
- **Jurkat** (ATCC TIB-152) – A human acute T-cell leukaemia line established from the peripheral blood of a 14-year-old male patient, commonly used in apoptosis signalling and immune cell cytotoxicity studies [2].
- **HUV-EC-C** (ATCC CRL-1730) – A human umbilical vein endothelial cell line isolated from the vein of the umbilical cord, used as a standard model for vascular biology and endothelial toxicity [3].
- **SH-SY5Y** (ATCC CRL-2266) – A thrice-subcloned human neuroblastoma cell line derived from the parental SK-N-SH line (ATCC HTB-11), established from a metastatic bone tumour of a 4-year-old female patient; widely used as an in vitro model of neuronal function and dopaminergic neurotoxicity [4].
- **BJ** (ATCC CRL-2522) – A normal human skin fibroblast line established from neonatal male foreskin tissue; used as a non-transformed somatic reference for cytotoxicity assessment [5].
- **MRC-5** (ATCC CCL-171) – A normal human lung fibroblast line derived from the lung tissue of a 14-week-old male embryo; a standard diploid reference line used to evaluate pulmonary cytotoxicity and genotoxicity [6].
- **Mesangial** – Mouse glomerular mesangial cells used to model renal function and nephrotoxicity in the kidney filtration unit [7].
- **SK-N-SH** (ATCC HTB-11) – A human neuroblastoma line established from the bone marrow aspirate of a 4-year-old female patient; used to investigate CNS cytotoxicity and neuronal differentiation [8].
- **H-4-II-E** (ATCC CRL-1548) – A rat hepatoma cell line used as a model of liver metabolism and hepatotoxicity in rodent systems, particularly for studying cytochrome P450-mediated bioactivation [9].
- **HEK293** (ATCC CRL-1573) – A human embryonic kidney cell line isolated from the kidney of a human embryo; widely used in toxicology as a cytotoxicity surrogate owing to its robust growth and high transfection efficiency [10].
- **N2a** (ATCC CCL-131) – A mouse neuroblastoma cell line (Neuro-2a) isolated from a spontaneous tumour of a strain A albino mouse; used as a model for neuronal differentiation and neurotoxicity [11].
- **NIH 3T3** (ATCC CRL-1658) – A mouse embryonic fibroblast line established from NIH/Swiss mouse embryo cultures; a standard non-tumourigenic reference used to benchmark cytotoxic responses [12].
- **Renal Proximal Tubule** (ATCC CRL-4031) – An hTERT-immortalised human renal proximal tubule epithelial cell line (RPTEC/TERT1) that closely recapitulates the functional properties of normal proximal tubular epithelium; the preferred in vitro model for evaluating direct nephrotoxicity [13].

### Graph Transformer Architectures

**Graphormer.** Atom feature vectors are projected to a 256-dimensional hidden space and augmented with learnable degree embeddings encoding local valence context. Four `TransformerConv` layers apply multi-head attention (4 heads, head dimension 64) with bond features as edge attributes and a learnable bias scalar ($\beta = \text{True}$), each followed by LayerNorm, GELU activation, and dropout ($p = 0.3$). Global mean pooling produces the graph-level embedding.

**GraphGPS.** Atom features are concatenated with an 8-dimensional Laplacian positional encoding (LPE) computed from the $k$ smallest non-trivial eigenvectors of the normalised graph Laplacian, then projected to 256 dimensions. Each of the four GPS layers applies `GINEConv` for local neighbourhood aggregation and multi-head self-attention (4 heads) over the full node set in parallel; outputs are combined via a residual LayerNorm with dropout ($p = 0.3$). Global mean pooling produces the graph-level embedding.

Both models share a two-layer MLP classification head ($256 \rightarrow 128 \rightarrow 1$) trained with class-ratio-weighted binary cross-entropy loss, the Adam optimiser, and a ReduceLROnPlateau scheduler (factor = 0.5, patience = 5). Early stopping (patience = 15, maximum 100 epochs) and a batch size of 64 were used consistently.

### Hyperparameter Settings

We adopted the Scikit-learn library's implementations of the three classical machine learning models. For Random Forest, 16 configurations were explored: `n_estimators` $\in$ {300, 500}, `max_features` $\in$ {sqrt, 0.3}, `min_samples_leaf` $\in$ {1, 3}, and `max_depth` $\in$ {None, 20}. `class_weight` was set to *balanced*. For SVM, nine configurations were evaluated over $C \in \{1, 10, 50\}$ and $\gamma \in \{\text{scale}, 0.01, 0.001\}$, with probability outputs enabled. For XGBoost, 12 configurations were tested spanning `n_estimators` $\in$ {300, 500}, `max_depth` $\in$ {4, 6, 8}, and `learning_rate` $\in$ {0.05, 0.1}; `subsample` and `colsample_bytree` were fixed at 0.8.

All GNN models were optimised using the Adam optimiser [14] with weight decay $1\times10^{-5}$. A ReduceLROnPlateau scheduler (factor = 0.5, patience = 10) was applied, and early stopping (patience = 20 epochs, maximum 100 epochs, batch size = 64) was used to prevent overfitting. The hyperparameter search covered hidden dimension $\in$ {128, 256}, message-passing layers $\in$ {3, 4}, dropout rate $\in$ {0.3, 0.4}, and learning rate $\in$ {0.0005, 0.001}.

Graph Transformer models were selected from 12 configurations: layers $\in$ {4, 6, 8}, hidden dimension $\in$ {256, 512}, attention heads = hidden/64, and dropout $\in$ {0.2, 0.3}. The search selected hidden 256, 4 layers, 4 heads, and dropout 0.3 for Graphormer, and hidden 256, 4 layers, 4 heads, and dropout 0.2 for GraphGPS. GraphGPS additionally used Laplacian positional encodings of dimension 8. Both models were trained with Adam and a ReduceLROnPlateau scheduler (factor = 0.5, patience = 5), with learning rate $10^{-3}$ in Stage 1 (pretraining on Proxy-DILI, up to 100 epochs, patience 20) and $5 \times 10^{-4}$ in Stage 2 (fine-tuning on the DILI train data, up to 80 epochs, patience 15). Class imbalance was handled via `BCEWithLogitsLoss` with `pos_weight` = 2.06 in Stage 1 and 1.85 in Stage 2. Graphormer used early stopping on validation ROC-AUC. GraphGPS used a combined score of $0.3 \times$ ROC-AUC + $0.7 \times$ F1, where F1 was evaluated at the threshold maximising F1 on the validation precision–recall curve; this threshold was carried forward to test time.

### Class Balancing: Undersampling Ratio Comparison

For the HepG2 caspase and CHO-K1 caspase datasets, an alternative undersampling ratio of 5:1 (inactive:active) was evaluated alongside the 1:1 scheme used in the main analysis. For the HepG2 caspase dataset, the 5:1 scheme yielded substantially inferior performance relative to the 1:1 scheme (consensus ROC-AUC of 0.835 vs. 0.915; MCC of 0.246 vs. 0.659), confirming that aggressive balancing was preferable even under conditions of extreme class asymmetry. A similar pattern was observed for the CHO-K1 caspase dataset, where the 1:1 scheme consistently outperformed the 5:1 alternative across all metrics at both decision thresholds.

### References


(1) ATCC Hep G2 [HEPG2] cell line (HB-8065), American Type Culture Collection, `https://www.atcc.org/products/hb-8065`.

(2) ATCC Jurkat, Clone E6-1 cell line (TIB-152), American Type Culture Collection, `https://www.atcc.org/products/tib-152`.

(3) ATCC HUV-EC-C [HUVEC] cell line (CRL-1730), American Type Culture Collection, `https://www.atcc.org/products/crl-1730`.

(4) ATCC SH-SY5Y cell line (CRL-2266), American Type Culture Collection, `https://www.atcc.org/products/crl-2266`.

(5) ATCC BJ cell line (CRL-2522), American Type Culture Collection, https://www.atcc.org/products/crl-2522.

(6) ATCC MRC-5 cell line (CCL-171), American Type Culture Collection, https://www.atcc.org/products/ccl-171.

(7) Xia, M.; Huang, R.; Witt, K. L.; Southall, N.; Fostel, J.; Cho, M.-H.; Jadhav, A.; Smith, C. S.; Inglese, J.; Portier, C. J.; Tice, R. R.; Austin, C. P. Compound cytotoxicity profiling using quantitative high-throughput screening. *Environmental Health Perspectives* **2008**, *116*, 284–291.

(8) ATCC SK-N-SH cell line (HTB-11), American Type Culture Collection, https://www.atcc.org/products/htb-11.

(9) ATCC H-4-II-E cell line (CRL-1548), American Type Culture Collection, https://www.atcc.org/products/crl-1548.

(10) ATCC 293 [HEK-293] cell line (CRL-1573), American Type Culture Collection, https://www.atcc.org/products/crl-1573.

(11) ATCC Neuro-2a cell line (CCL-131), American Type Culture Collection, https://www.atcc.org/products/ccl-131.

(12) ATCC NIH/3T3 cell line (CRL-1658), American Type Culture Collection, https://www.atcc.org/products/crl-1658.

(13) ATCC RPTEC/TERT1 cell line (CRL-4031), American Type Culture Collection, https://www.atcc.org/products/crl-4031.

(14) Kingma, D. P.; Ba, J. Adam: A method for stochastic optimization. *arXiv preprint arXiv:1412.6980* **2014**.
